# *Cis*-regulatory variation in *ISOCHORISMATE SYNTHASE 1* modulates systemic salicylic acid biosynthesis and systemic acquired resistance in plants

**DOI:** 10.1101/2025.07.17.664945

**Authors:** Jiyeon Hyun, Rabia Ahuja, Hunter David Rose, Mariana Lico De Freitas, Haley Welker, Alyssa Marie Curtis, Amanda Navodani Hewa Maithreege, Chan Yul Yoo, Heejin Yoo

**Author notes:** These authors contributed equally to this work (Jiyeon Hyun, Rabia Ahuja). Corresponding author: Dr. Heejin Yoo Biology Department University of Utah, Salt Lake City, Utah, 84112 USA.

## Abstract

Systemic acquired resistance (SAR) is a long-distance immune response that protects uninfected plant tissues following a localized pathogen attack. In the model plant *Arabidopsis thaliana*, SAR depends on the systemic accumulation of salicylic acid (SA), mediated by the transcription factor CCA1 HIKING EXPEDITION (CHE), which activates the SA biosynthetic gene *ISOCHORISMATE SYNTHASE1* (*ICS1*). However, the conservation and functional significance of the CHE-*ICS1* regulatory module across plant species remain poorly understood, particularly in the Brassicaceae, where ICS1 is the major contributor to SA biosynthesis among two known pathways: the ICS1 and PHENYLALANINE AMMONIA-LYASE (PAL) routes. Here, we identify natural variation in *cis*-regulatory elements within the *ICS1* promoter that affects CHE binding across species. In multiple Brassicaceae species with divergent *cis*-element sequences, CHE fails to regulate *ICS1*, leading to the absence of systemic *ICS1* induction and SA accumulation following pathogen infection. Despite this deficiency, SAR still occurs in these species, albeit to a lesser extent than in species with successful systemic induction of SA mediated by an intact CHE-*ICS1* regulatory module. Interestingly, introducing the CHE-*ICS1* module into species lacking this interaction confers systemic SA accumulation, highlighting the potential to enhance systemic immunity through *cis*-element modification. Our findings demonstrate that sequence variation in a *cis*-regulatory element underlies the interspecies diversification of SAR regulatory mechanisms and highlight the evolutionary plasticity of plant immune signaling. This study provides a molecular framework for engineering enhanced systemic immunity in crops, particularly within the Brassicaceae, through the targeted modification of *cis*-regulatory elements that regulate SA biosynthesis.

## Introduction

Plant pathogens cause substantial yield losses and pose a serious threat to global food security. To combat infection, plants have evolved a multilayered immune system that includes the production of defense-related phytohormones such as salicylic acid (SA). SA is a key signaling molecule that is essential for immunity against biotrophic and hemibiotrophic pathogens, and its accumulation is tightly regulated to balance defense and growth (Vlot et al., 2009; Fu and Dong, 2013; Peng et al., 2021; Ullah et al., 2023). The precise temporal and spatial regulation of SA biosynthesis is crucial, as constitutive synthesis can lead to fitness penalties due to the growth– defense tradeoff (Bowling et al., 1997; Ding and Ding, 2020). In *Arabidopsis thaliana*, endogenous SA levels oscillate in a circadian pattern, peaking at dawn to anticipate pathogen attack (Wang et al., 2011; Goodspeed et al., 2012; Zheng et al., 2015; Karapetyan & Dong, 2018). Upon pathogen infection, SA accumulates substantially at the site of infection to activate local defense responses, and it accumulates to a lesser extent in uninfected systemic tissue to establish systemic acquired resistance (SAR), a long-lasting, broad-spectrum immune response that primes uninfected distal tissue to respond more rapidly and robustly to subsequent pathogen challenge (Fu and Dong, 2013; Ding and Ding, 2020; Zheng et al., 2015).

SA biosynthesis occurs via two distinct pathways: the isochorismate synthase 1 (ICS1)-mediated pathway and the phenylalanine ammonia-lyase (PAL)-mediated pathway (Wildermuth et al., 2001). The ICS1 pathway involves the conversion of chorismate to isochorismate in plastids, followed by cytosolic steps that are mediated by EDS5 and then PBS3, followed by either EPS1 or spontaneous conversion (Rekhter et al., 2019; Torrens-Spence et al., 2019). In contrast, the PAL pathway converts phenylalanine to cinnamic acid via PAL and ultimately to SA through intermediate steps that involve AIM1 and other enzymes, many of which remain unidentified or incompletely characterized (Jia et al., 2023; Ullah et al., 2023). Although both routes are functionally active, the predominant route in each plant species and the regulatory logic behind this selection remain poorly understood and are not well characterized.

The ICS1 pathway is generally considered the primary route in Arabidopsis and other Brassicaceae species, whereas the PAL pathway appears to dominate in other lineages, including monocots and early-diverging taxa (Hong et al., 2025; Jia et al., 2023; Torrens-Spence et al., 2024).

This pattern aligns with the broader conservation of PAL-pathway genes, such as *PAL* and *AIM1*, across land plants, whereas functional ICS1 pathway components, including *PBS3* and *EPS1*, are restricted to the Brassicales. While this trend suggests that SA biosynthesis strategies vary across lineages, the specific pathways contributing to SA accumulation in different tissue types or biological contexts remain relatively unclear. In particular, the process of SA synthesis in uninfected systemic tissue during SAR is still poorly characterized in most plant species.

In Arabidopsis, systemic SA accumulation relies largely on the transcription factor CCA1 HIKING EXPEDITION (CHE), a member of the plant-specific TCP family (Pruneda-Paz et al., 2009; Zheng et al., 2015). Notably, SA biosynthesis in locally infected and systemic tissues is regulated by distinct mechanisms. While CHE is not essential for local SA production, it is required for systemic SA induction following local pathogen attack, as well as for maintaining the circadian oscillation of SA levels in Arabidopsis (Zheng et al., 2015). CHE binds to a conserved TCP motif (5’-GTGGGCCC-3’) in the *ICS1* promoter to activate *ICS1* transcription and promote SA accumulation in systemic tissues (Zheng et al., 2015). While other transcription factors SARD1 and CBP60g also contribute to systemic SA accumulation, the complete loss of systemic SA induction in the *che-2* mutant, along with the significantly reduced expression of *SARD1* and *CBP60g* in this background, suggests that CHE functions as an upstream and central regulator in the systemic SA signaling cascade (Zheng et al., 2015).

Recent evidence further highlights CHE’s role in linking local infection to systemic immunity in Arabidopsis. In systemic tissues, CHE undergoes sulfenylation in response to hydrogen peroxide (H_2_O_2_), a mobile signal generated from local infection sites (Cao et al., 2024). This redox modification enhances CHE’s binding affinity for the *ICS1* promoter, thereby promoting systemic SA biosynthesis. In addition to its direct impact on *ICS1*, CHE also contributes to the regulation of other key SAR components, including pipecolic acid (Pip) and N-hydroxy-pipecolic acid (NHP) (Bernsdorff et al., 2016; Hartmann et al., 2018; Yildiz et al., 2021), highlighting its expansive role in orchestrating systemic immunity (Cao et al., 2024).

Although the CHE-*ICS1* regulatory module has a well-established role in systemic SA regulation in Arabidopsis, it remains largely unknown whether it is conserved or functional in other plant species, particularly within the Brassicaceae, where ICS1 is known to be the major route of SA biosynthesis. The biological functions of CHE orthologs in non-Arabidopsis species have been explored only minimally, and a substantial research gap remains to determine whether the *ICS1* promoters of other species retain the TCP-binding site necessary for the CHE-mediated activation of *ICS1* during SAR.

In this study, we investigate the evolutionary and functional diversity of the CHE–*ICS1* regulatory module across plant species, including the genetic variation in CHE proteins and TCP-binding motifs in *ICS1* promoters. By assessing the impacts of the genetic variation on CHE binding, *ICS1* transcription, and systemic SA accumulation, we uncover how *cis*-element variants influence the regulation of SA, specifically in distal, uninfected tissues. We demonstrate that a single-nucleotide substitution within the TCP motif of the *ICS1* promoter is sufficient to disrupt the CHE-*ICS1* interaction and impair both systemic *ICS1* transcription and SA accumulation. Interestingly, species lacking systemic SA accumulation are still able to mount SAR, albeit to a limited extent. Notably, introducing the CHE–*ICS1* module into *B. napus*, which naturally lacks it, confers the capacity for systemic SA accumulation. This work provides critical mechanistic insights into how natural variation in *cis*-elements can modulate a core SA biosynthetic pathway, thereby shaping the capacity for SA-based systemic immune signaling. Our findings advance the current understanding of species-specific immune regulation and underscore the evolutionary impact of *cis*-regulatory variation on plant defense.

## Results

### Evolutionary origin and conservation of CHE in plants

CHE is a member of the TEOSINTE BRANCHED1/CYCLOIDEA/PCF (TCP) transcription factor family, which is defined by a conserved, non-canonical basic helix-loop-helix (bHLH) DNA-binding TCP domain in the N-terminus. The TCP family has undergone extensive expansion through multiple rounds of gene duplication, subfunctionalization, and neofunctionalization during evolution (Navaud et al., 2007). To investigate when CHE diverged from other TCP family members and identify its orthologs across plant lineages, we performed a comprehensive phylogenetic analysis of TCP transcription factors using representative genomes from 52 species including 1 rhodophyte, 10 chlorophytes, 1 charophyte, 1 hornwort, 1 liverwort, 3 mosses, 1 lycophyte, 1 fern, 2 gymnosperms, and 31 angiosperms. A total of 968 TCP genes were identified across 40 species via a homology search using the TCP domain sequences of Arabidopsis CHE (TCP21, Class I) and TCP1 (Class II) as queries (**Supplementary Table 1**). A maximum-likelihood tree constructed using full-length protein sequences resolved the TCP family into 12 distinct clades (Clades A–L, **Figure 1A)**.

**Figure 1.**
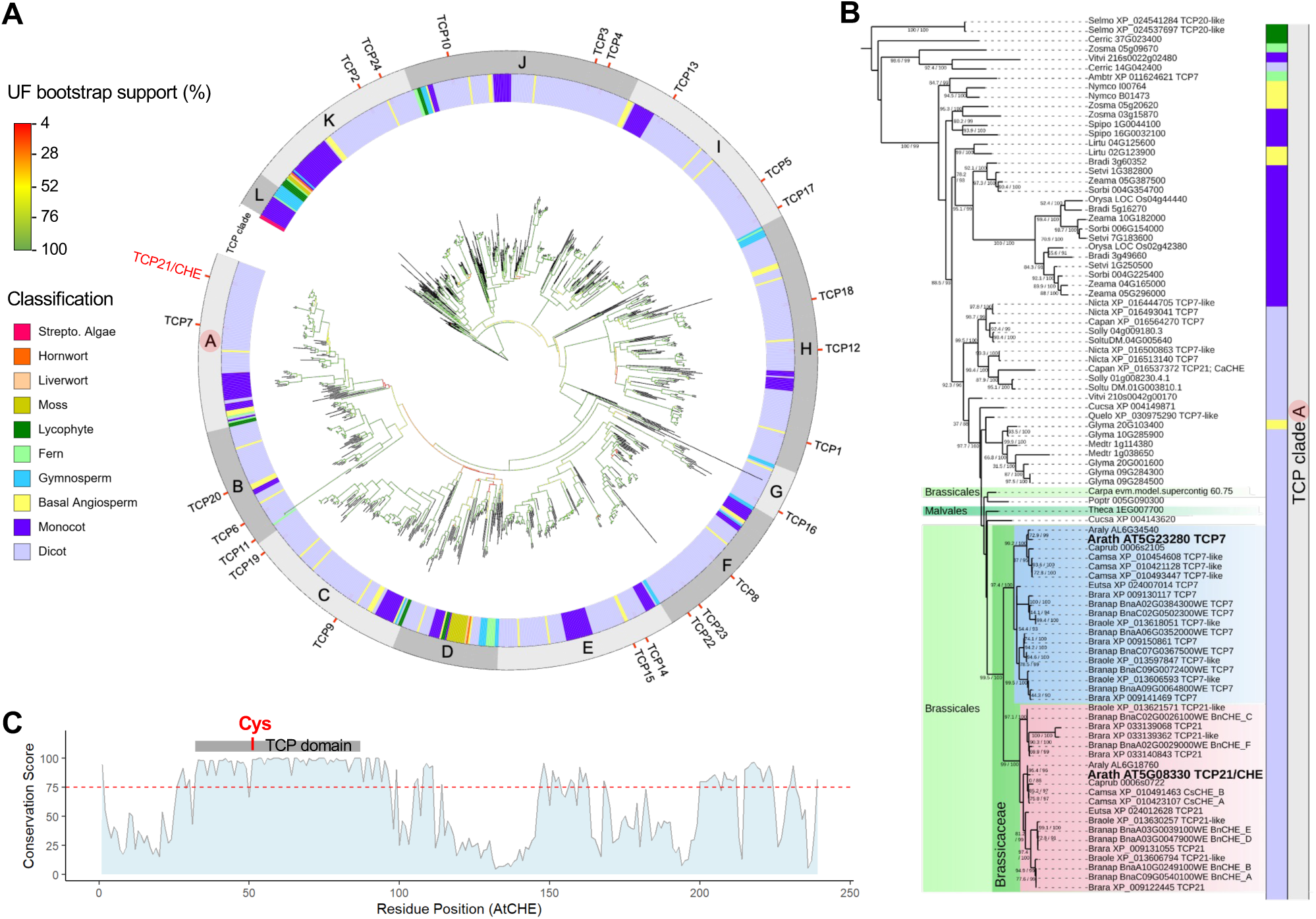
Phylogenetic and sequence analyses of the TCP transcription factor family across the plant kingdom. **(A)** Phylogenetic tree of the TCP family. The color-coded strip around the tree indicates the taxonomic group. TCP proteins are classified into 12 clades based on tree topology and are marked in the adjacent gray strip. The TCPs from *Arabidopsis thaliana* (TCP1-24) are labeled. CHE (TCP21) from *A. thaliana* is highlighted in red in the tree. **(B)** Phylogenetic tree of Clade A TCPs, which include Arabidopsis CHE. **(C)** Residue-level conservation of Clade A TCPs. Conservation scores for each residue are mapped onto the CHE protein sequence from *A. thaliana*. The TCP domain, as well as the conserved cysteine (Cys) residue proposed to undergo sulfenylation, are indicated.

Arabidopsis CHE, alongside its paralog TCP7 (At5g43280), clusters in Clade A. Clade A TCPs are conserved across dicots, monocots, basal angiosperms, lycophytes, and ferns, but were not detected in the surveyed gymnosperms, suggesting an evolutionary loss of Clade A TCPs in that lineage. Clade A TCPs further diverged into two sub-clades each including Arabidopsis TCP7 and CHE (**Figure 1B**). A phylogenetic comparison indicates that this divergence likely occurred after the emergence of the Brassicales order but prior to the diversification of the Brassicaceae family, since both sub-clades are conserved across Brassicaceae species, but are absent in non-Brassicaceae Brassicales (e.g., *Carica papaya*) and in the Malvales (e.g., *Theobroma cacao*), the closest outgroup to the Brassicales (**Figure 1B**). Outside the Brassicaceae, the Clade A TCPs are the most closely related orthologs of Arabidopsis CHE, spanning lineages from lycophytes to angiosperms. These TCPs may represent ancestral forms from which both CHE and TCP7 evolved, or they could be independently derived proteins that retain key ancestral features shared with CHE.

To assess the conservation of the Clade A TCPs, we calculated residue-level conservation scores from a multiple sequence alignment of the TCPs. These scores reflect the percentage of TCPs sharing the most common amino acid at each alignment position. Mapping these values onto the Arabidopsis CHE sequence revealed that over 51% of residues are conserved in more than 75% of Clade A TCPs. The TCP domain, which is critical for DNA binding, is particularly well conserved, with an average conservation score of 96.4%, suggesting the functional conservation of DNA-binding activity across Clade A TCPs (**Figure 1C**). Notably, a cysteine residue, previously shown to undergo sulfenylation in systemic tissue following local pathogen infection to enhance CHE binding to the *ICS1* promoter (Cao et al., 2024), was conserved across all tested Clade A TCPs except one from *Vitis vinifera*. This observation suggests a redox-sensitive regulatory mechanism that is conserved across plant lineages, from early vascular plants to flowering plants, supporting a potential role for Clade A TCPs in the regulation of SAR.

### Evolutionary diversity of TCP *cis*-regulatory elements in *ICS1* promoters across plant species

Beyond the conservation of its functions, CHE’s regulatory interactions with *ICS1* are likely shaped by variation in *cis*-regulatory elements. We therefore examined whether the CHE-binding motif is conserved in *ICS1* promoters across a variety of species. We analyzed the 2-kb upstream regions of *ICS1* orthologs from a broad range of flowering plants, including basal angiosperms, monocots, and dicots. Promoter sequences were scanned for CHE-binding motifs using the FIMO tool in MEME Suite 5. 5. 7 (Charles et al, 2011). The canonical CHE binding motif (GTGGGnCC), which is highly conserved among Class I TCP proteins, was obtained from the Plant Transcription Factor Database (Tian et al., 2020) and used as the primary query. In addition, the motif search was extended to include three nucleotides downstream of the canonical site (GTGGGnCC<u>NNN</u>), as recent structural studies of TCP–DNA complexes have proposed a three-site recognition mode that includes the adjacent downstream CAC trinucleotide (GTGGGnCC<u>CAC</u>; Zhang et al., 2023).

The analysis revealed widespread variation in both the sequence and position of the TCP-binding sites across *ICS1* promoters (**Figure 2**). While the motif’s position in the promoter is highly conserved within the Brassicaceae, other families exhibit greater variability in both motif sequence and position. Within the Brassicaceae, two distinct motif variants are identified corresponding to the two major lineages: Linage I (Arabidopsis, *Camelina sativa*, *Capsella rubella*, and *Arabidopsis lyrata*) and Lineage II (*Brassica rapa*, *B. oleracea*, *B. napus,* and *Eutrema salsugineum*). Lineage I species retain a highly conserved TCP-binding motif nearly identical to that of Arabidopsis, including conserved C(A/T)G downstream trinucleotides. In contrast, Lineage II species share a G-to-A substitution at the first position of the motif, along with greater variation in the three downstream nucleotides, raising the question of whether these changes alter the CHE-mediated regulation of *ICS1*. Similar motif variations were found in several Solanaceae species, including *Solanum tuberosum*, *S. lycopersicum*, and *Capsicum annuum*, which also exhibit G-to-A substitutions and divergence in their downstream trinucleotides. In monocots, the TCP-binding motifs were more divergent and displayed greater positional variability compared to dicots, suggesting a distinct regulatory architecture for *ICS1* expression. Taken together, this comparative analysis reveals natural variation in both the sequence and positioning of the TCP-binding motif in *ICS* promoters.

**Figure 2.**
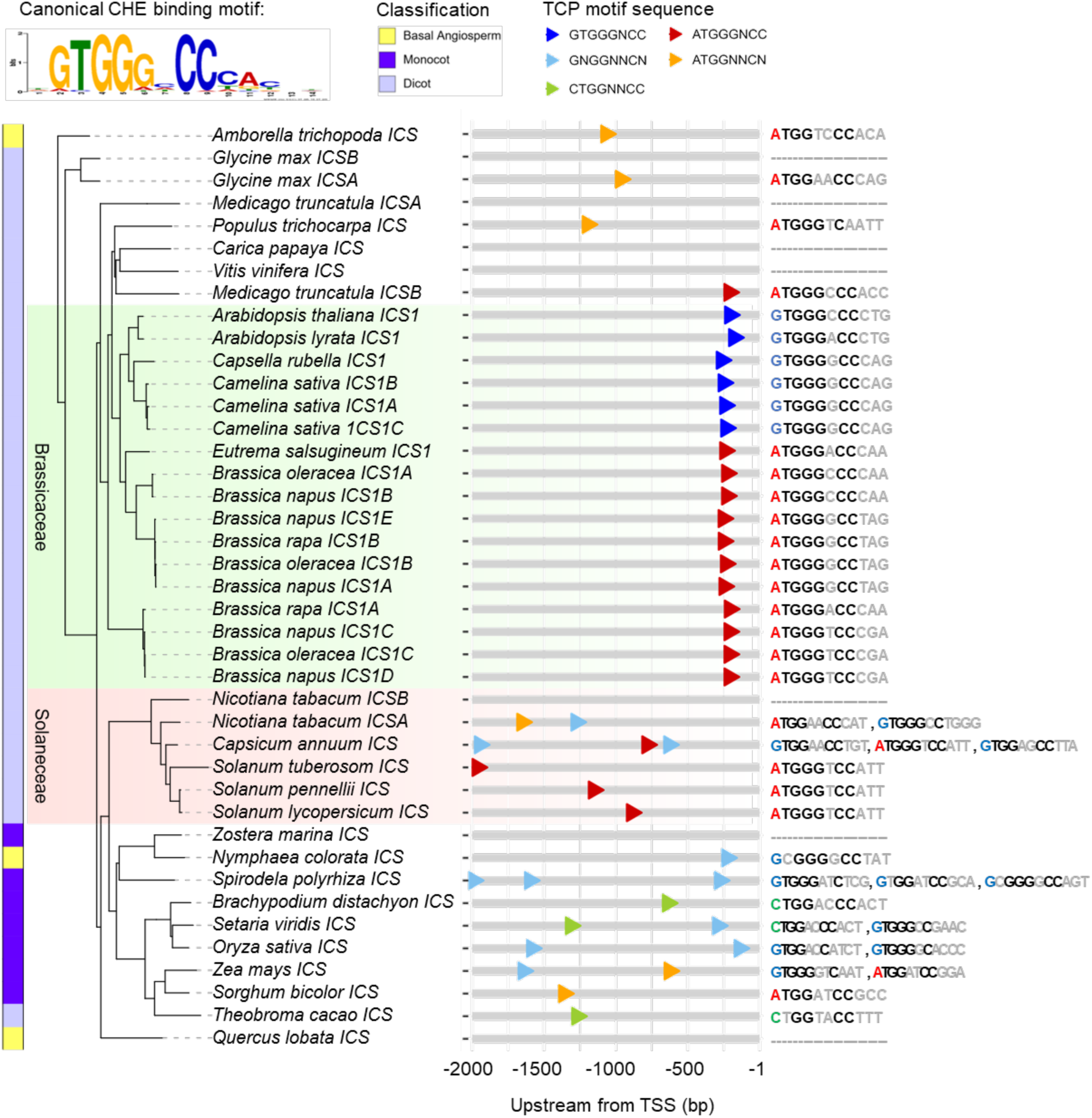
Comparative analysis of TCP-binding motifs in *ICS1* promoters across plant species. The distribution and sequence variation of predicted TCP-binding motifs within *ICS* promoter regions spanning 2 kb upstream of the coding sequence are shown. Promoter regions are represented as gray bars, and predicted TCP binding motifs are indicated by colored arrowheads, with colors denoting sequence variants. The phylogenetic relationships of *ICS* orthologs are shown on the left, and the TCP-binding motif sequences are listed on the right.

### *Cis*-element variation drives species-specific CHE binding to the *ICS1* promoter in a yeast one-hybrid system

To investigate whether the variations in TCP *cis*-elements within *ICS* promoters affect CHE’s binding to *ICS1* promoters, we performed yeast one-hybrid (Y1H) assays using CHE orthologs (Clade A TCP or CHE-like subclade members) and *ICS1* promoter constructs from multiple species. We selected three Brassicaceae species: Arabidopsis, *C. sativa*, and *B. napus*. *C. sativa* retains a TCP-binding site highly conserved with Arabidopsis, while *B. napus* carries a G-to-A substitution in this motif.

All tested CHE orthologs, including AtCHE (Arabidopsis), CsCHE_A (*C. sativa*), and BnCHE_D (*B. napus*), could bind the *ICS1* promoters from *A. thaliana* and *C. sativa*, except BnCHE_F (**Figure 3A**). Additional *B. napus* CHE paralogs BnCHE_A, BnCHE_B, BnCHE_C, and BnCHE_E also bound the Arabidopsis *ICS1* promoter **(Supplementary Figure 1A**). However, none of the CHE proteins bound the *B. napus ICS1* promoters (**Figure 3A; Supplementary Figure 1B-E**), indicating that *cis*-element divergence in *B. napus* impairs CHE recruitment.

**Figure 3.**
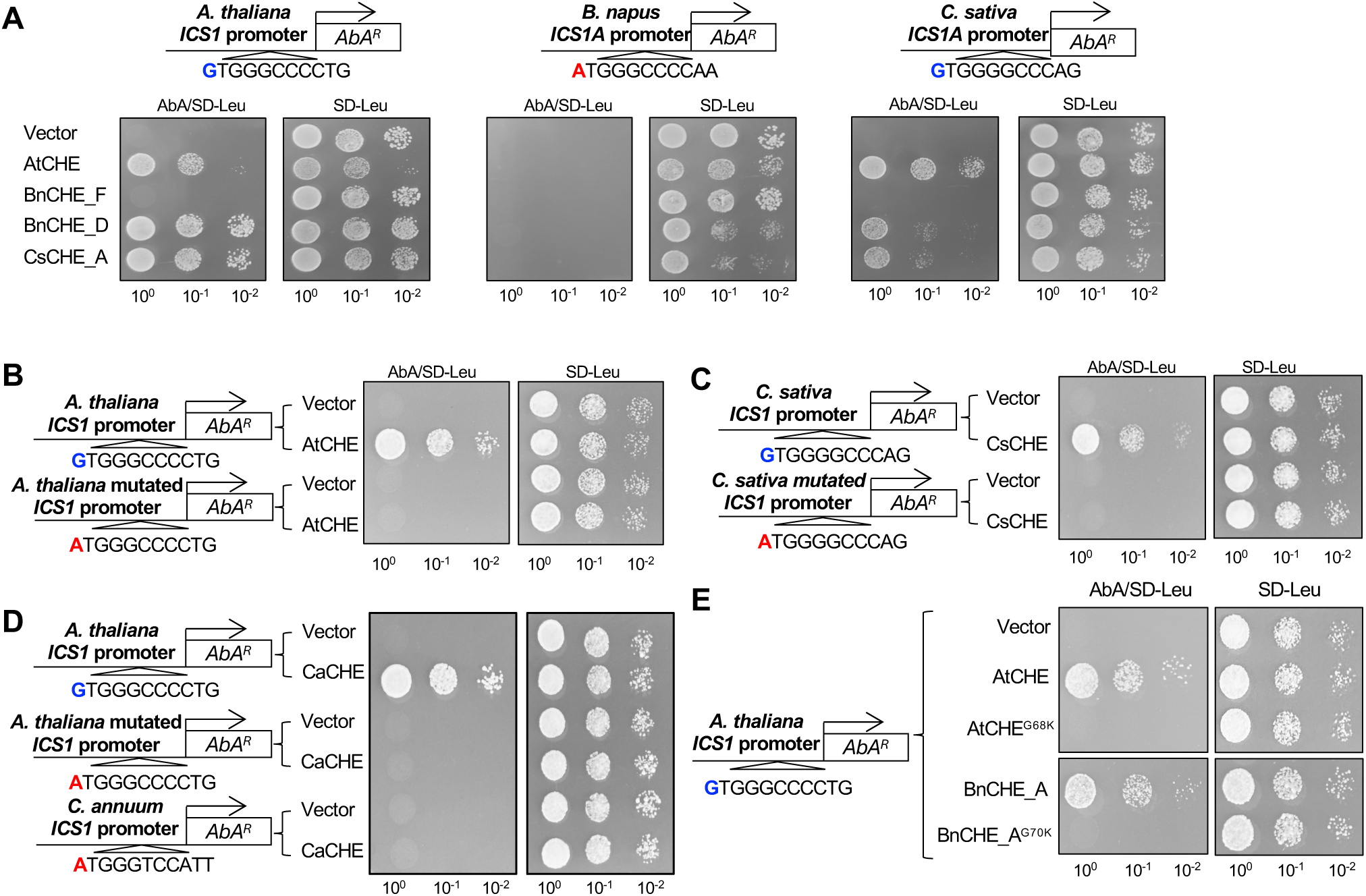
Variations in the *cis*-regulatory motif of the *ICS1* promoter and in the TCP domain of CHE affect CHE binding to the *ICS1* promoter. Yeast one-hybrid (Y1H) assays testing CHE binding to *ICS1* promoters from multiple species. **(A)** The interactions between CHE proteins and *ICS1* promoters from *A. thaliana*, *C. sativa*, and *B. napus* are shown. Arabidopsis CHE (AtCHE), *B. napus* CHEs (BnCHE_F and BnCHE_D), and *C. sativa* CHE (CsCHE_A) were tested for their interactions with *AtICS1*, *BnICS1_A*, and *CsICS1_A.* (B–C) Mutation of the conserved G to an A in the TCP motifs of *A. thaliana* (B) and *C. sativa* (C) abolishes CHE binding. (D) *C. annuum* CHE (CaCHE) interacts with *AtICS1* but not with *CaICS1*, which contains a noncanonical TCP motif (ATGGGTCATT) or with G-to-A mutated *AtICS1*. **(E)** CHEs containing a lysine instead of a glycine in the TCP domain do not bind to *ICS1*. The mutation of the glycine to lysine in AtCHE (AtCHE – G68K) abolishes binding to *AtICS1*, mirroring BnCHE_F. The same mutation in BnCHE_A also abolishes binding to *AtICS1*.

To directly test the functional importance of the conserved guanine at the first position of the canonical TCP-binding site, we introduced a single G-to-A substitution into the *A. thaliana* and *C. sativa ICS1* promoters. This single nucleotide change was sufficient to disrupt the CHE-*ICS1* interaction, as neither AtCHE nor CsCHE_A could bind the G-to-A mutated promoters (**Figure 3B** and **3C**), confirming that the first nucleotide in the TCP-binding site is critical for CHE recruitment. We then expanded our interaction tests to another species, *C. annuum* from the Solanaceae, which shares the same G-to-A substitution as *B. napus* and has two additional non-canonical TCP motifs that retain the core guanine at the first position but vary at other positions (**Figure 2**). Similar to *B. napus* CHE, while *C. annuum* CHE (CaCHE) could bind the Arabidopsis *ICS1* promoter, it failed to interact with both the native *C. annuum ICS1* promoter, which contains divergent TCP-binding sites, and with the G-to-A mutated Arabidopsis *ICS1* promoter (**Figure 3D**).

Unlike most CHE orthologs tested, BnCHE_F was unable to bind any of the tested *ICS1* promoters regardless of whether the promoters contained a canonical TCP-binding motif (**Figure 3A**). We further investigated the binding ability of CHE orthologs from *B. rapa* and *B. oleracea*, the parental species of *B. napus*, and found that while all BoCHEs bound the *AtICS1* promoter, some BrCHEs could not (**Supplementary Figure 1F**). Notably, all CHE proteins that failed to bind to the *ICS1* promoter shared a glycine-to-lysine (G→K) substitution within the TCP domain (**Supplementary Figure 1G**), implicating the importance of this residue in DNA-binding capacity. To directly test whether this G→K substitution disrupts CHE-*ICS1* interaction, we introduced the same G→K mutation into AtCHE and BnCHE_A, both of which are capable of binding the canonical TCP-binding site. Strikingly, the G→K mutants failed to bind the *AtICS1* promoter, similar to BnCHE_F (**Figure 3F**), confirming that the conserved glycine residue is essential for CHE–DNA interactions.

Together, these findings demonstrate that both *cis*-element variation in the *ICS1* promoter and amino acid substitutions in the TCP domain of CHE proteins can independently disrupt the CHE–*ICS1* interaction and contribute to regulatory divergence between species.

### *In planta* validation of species-specific CHE-*ICS1* promoter interactions

To assess the impact of TCP-binding site variation on the CHE-*ICS1* interaction *in planta*, we first conducted transient GUS expression assays using Arabidopsis protoplasts isolated from both wild-type Col-0 and *che-2*. To examine the CHE-*ICS1* interaction during SAR, leaves were infected with *P. syringae* pv. *maculicola* (*Psm*) ES4326 carrying the *AvrRpt2* effector (*Psm* ES4326/*AvrRpt2*), and protoplasts were isolated from distal leaves 2 days after local infection. The protoplasts were transfected with GUS reporter constructs (**Figure 4A**) driven by the *ICS1* promoter from Arabidopsis, *C. sativa*, or *B. napus*, or a mutated Arabidopsis *ICS1* promoter carrying a G-to-A substitution at the first position of TCP-binding site. In *che-2* protoplasts, which have very low *CHE* levels due to a knock down mutation (Pruneda-Paz et al., 2009), AtCHE was co-transfected with each reporter construct to provide CHE activity.

**Figure 4.**
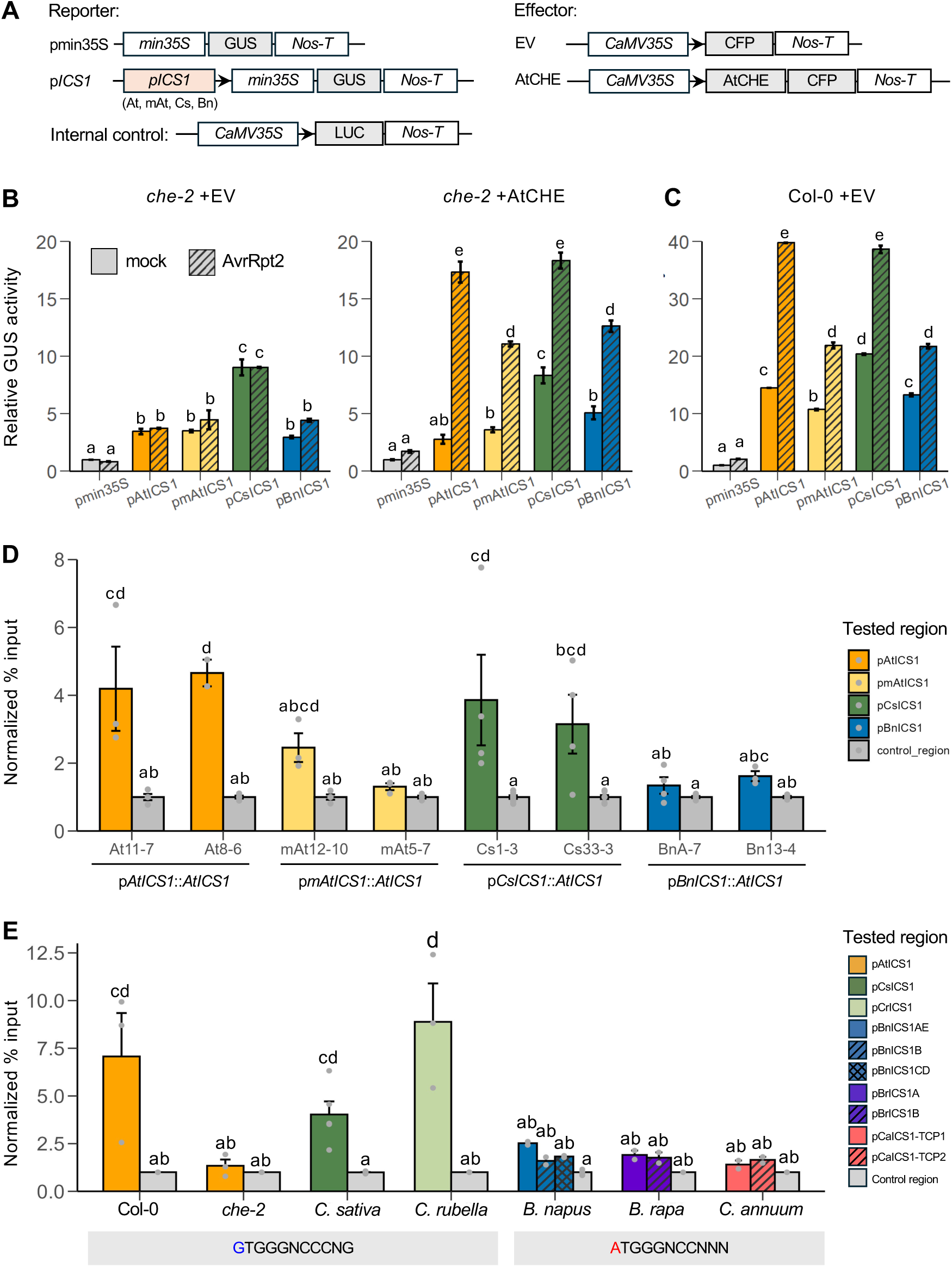
TCP-binding site variation affects CHE binding to *ICS1* promoters *in planta*. **(A)** Schematic of protoplast reporter constructs containing GUS driven by *ICS1* promoters from *A. thaliana*, *C. sativa*, and *B. napus*, as well as a G-to-A mutated *A. thaliana ICS1* promoter, and effector constructs expressing AtCHE or an empty vector (EV). Luciferase driven by the CaMV35S promoter was co-transfected as an internal control. **(B-C)** GUS reporter activity in protoplasts isolated from the systemic tissue of *che-2* (B) and Col-0 (C) after transfection with combinations of the reporter constructs described in (A), followed by local infection with *Psm* ES4326/*AvrRpt2*. GUS activity was normalized to luciferase activity from the co-transfected internal control. Values were further normalized to the pMIN35S control (set to 1) and are shown as relative GUS activity. Robust GUS activation was observed with the *AtICS1* and *CsICS1_A* promoters in the presence of AtCHE, especially following pathogen infection in *che-2*. In contrast, the *BnICS1* promoter and the G-to-A-mutated *AtICS1* promoter exhibited only modest induction. Error bars represent the SE. Different letters indicate statistically significant differences in relative GUS activity (*p* < 0.05, one-way ANOVA followed by Tukey’s multiple comparisons test). **(D)** Chromatin immunoprecipitation (ChIP)-qPCR with a polyclonal anti-AtCHE antibody reveals *in vivo* CHE binding to the *ICS1* promoter in transgenic Arabidopsis plants carrying one of various *ICS1* promoters. CHE bound more strongly to the *AtICS1* and *CsICS1* promoters than to a G-to-A mutated *AtICS1* promoter or the *BnICS1* promoter. Error bars represent the SE. Different letters indicate statistically significant differences in CHE binding to the tested DNA regions (*p* < 0.05, one-way ANOVA followed by Tukey’s multiple comparisons test). **(E)** ChIP-qPCR with an anti-AtCHE antibody reveals *in vivo* CHE binding to the *ICS1* promoter in various species. CHE bound strongly to *ICS1* promoters in *A. thaliana, C. sativa*, and *C. rubella*—all containing the canonical TCP-binding motif—but weakly in *B. napus, B. rapa, B. oleracea*, and *C. annuum*, whose promoters carry divergent TCP-binding site variants. Error bars represent the SE. Different letters indicate statistically significant differences in CHE binding to the tested DNA regions across species (*p* < 0.05, one-way ANOVA followed by Tukey’s multiple comparisons test).

In *che-2* protoplasts, when AtCHE was not co-transfected, no increase in promoter activity was observed following local pathogen infection, as reflected by GUS activity (**Figure 4B**). When AtCHE was co-transfected, however, all *ICS1* promoters exhibited stronger activity after local pathogen infection, confirming that CHE is required for *ICS1* activation during SAR. Notably, the *AtICS1* and *CsICS1* promoters exhibited stronger activation than the *BnICS1* and mutated *AtICS1* promoters, indicating the importance of the canonical TCP-binding site for the CHE-mediated activation of *ICS1* during SAR. A similar pattern was observed in Col-0 protoplasts, where the *AtICS1* and *CsICS1* promoters were more active than the *BnICS1* and *mAtICS1* promoters, confirming the importance of the canonical TCP-binding site for *ICS1* induction (**Figure 4C**).

To extend these findings *in planta*, we examined CHE-*ICS1* binding using Arabidopsis *salicylic acid induction-deficient 2* (*sid2*) mutants — which lack functional *ICS1* — by transgenically expressing the *ICS1* coding sequence under the control of one of four promoters: (1) the native *AtICS1* promoter driving the *AtICS1* coding sequence (p*AtICS1*::*AtICS1*); (2) a mutated *AtICS1* promoter (with a G-to-A substitution at the TCP-binding site) driving the *AtICS1* coding sequence (pm*AtICS1*::*AtICS1*) (3) the *CsICS1A* promoter driving the *CsICS1A* coding sequence (p*CsICS1*::CsICS1); and (4) the *BnICS1A* promoter driving the *BnICS1A* coding sequence (p*BnICS1*::*BnICS1*). Chromatin immunoprecipitation (ChIP) assays were conducted using a polyclonal anti-AtCHE antibody to assess CHE binding to each promoter (**Figure 4D**). Consistent with the protoplast-based assays, strong AtCHE binding was observed with the *AtICS1* and *CsICS1* promoters, whereas binding to the *mAtICS1* and *BnICS1* promoters was significantly lower, supporting the importance of the conserved TCP-binding site for CHE binding.

To further validate CHE–*ICS1* binding across species, we performed ChIP assays in multiple plant species using the anti-AtCHE antibody, which cross-reacts with CHE orthologs due to their high sequence conservation (**Figure 1**). Immunoblot analysis confirmed expression of CHE orthologs in tested species (**Supplementary Figure 2**). In addition to the species used in our Y1H assays, we included *C. rubella* and *B. rapa*. ChIP analysis revealed strong CHE binding to the *ICS1* promoters in *A. thaliana, C. sativa*, and *C. rubella*, all of which contain a conserved TCP-binding motif (**Figure 4E**). In contrast, CHE binding was substantially weaker in *B. napus, B. rapa*, and *C. annuum*, consistent with the sequence divergence in their TCP-binding regions. These findings demonstrate that *in vivo*, the CHE–*ICS1* interaction requires a conserved TCP-binding site.

### *ICS1* expression and SA levels in systemic tissues correlate with the CHE–*ICS1* interaction

To determine whether divergence in the TCP-binding site of the *ICS1* promoter compromises systemic SA induction along with CHE-*ICS1* interaction, we compared *ICS1* transcript levels and SA accumulation in two representative Brassicaceae species—*C. sativa* cv. Celine and *B. napus cv.* Westar—following infection with *Psm* ES4326/*AvrRpt2.* In locally infected tissues, both Brassicaceae species exhibited strong *ICS1* induction and robust SA accumulation by 1 dpi, indicating conserved local SA activation via the ICS1 pathway (**Figure 5**). However, in systemic tissue, only *C. sativa*, which harbors a canonical TCP-binding motif in the *ICS1* promoter that supports CHE-mediated regulation, showed significant *ICS1* upregulation and elevated SA levels. In contrast, *B. napus*, which carries a divergent TCP motif and lacks a detectable CHE-*ICS1* regulatory interaction, failed to induce *ICS1* or accumulate SA in systemic tissue throughout the tested time course. The pattern was consistent across different cultivars of each species, including *C. sativa* cv. Suneson and *B. napus* cv. Golden, as well as different species including *B. rapa* cv. K956 and *B. oleracea* cv. Crops-HH1, the two parental species of *B. napus* (**Supplementary Figure 3**). These findings demonstrate that systemic *ICS1* activation and SA biosynthesis correlate with the presence of a conserved TCP-binding site that facilitates CHE–*ICS1* interaction, suggesting that *cis*-regulatory divergence in the TCP-binding motif of the *ICS1* promoter underlies species-specific differences in systemic immune signaling.

**Figure 5.**
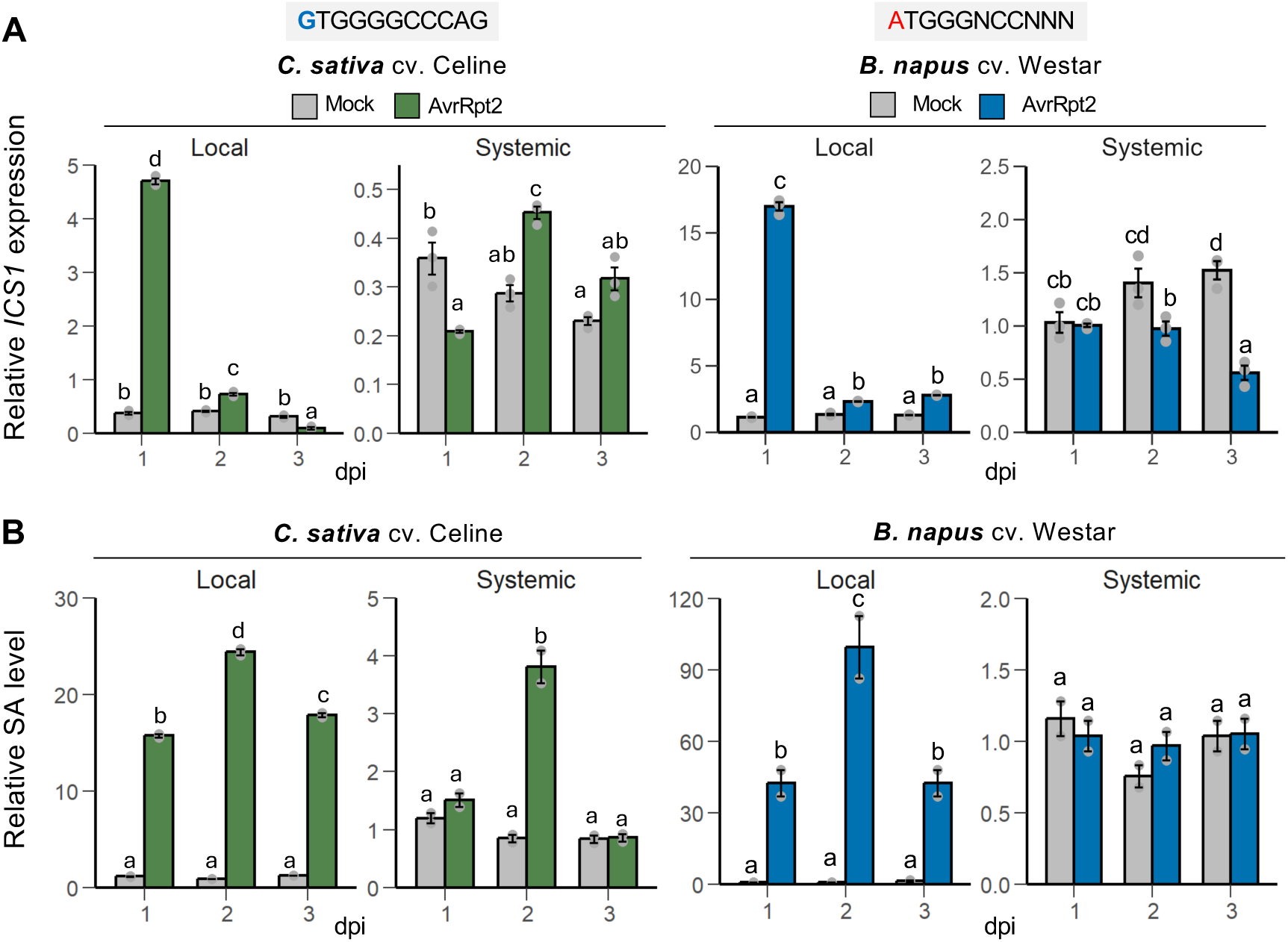
*ICS1* expression and SA accumulation in local and systemic tissue differ between *C. sativa* and *B. napus*. Systemic *ICS1* expression and SA accumulation were observed only *in C. sativa*; *B. napus* showed no systemic response. **(A–B)** Relative *ICS1* expression **(A)** and SA levels **(B)** were measured in local and systemic leaves from *C. sativa* cv. Celine and *B. napus* cv. Westar following infection with *Psm* ES4326/*AvrRpt2* (OD₆₀₀ = 0.02) or 10 mM MgCl₂ (mock) treatment at 1, 2, and 3 days post-infection (dpi). Both species showed strong *ICS1* induction and SA accumulation in locally infected tissue. However, only *C. sativa*, which retains the canonical TCP-binding site (GTGGGCCCAG) in the *ICS1* promoter, exhibited robust *ICS1* expression and SA accumulation in systemic tissue. In contrast, *B. napus*, which has a divergent TCP-binding motif (ATGGGNCCNNN), showed no systemic *ICS1* induction or SA accumulation. Error bars represent the SE. Different letters indicate statistically significant differences (*p* < 0.05, one-way ANOVA followed by Tukey’s multiple comparisons test).

### Plants lacking the CHE-*ICS1* interaction can only modestly induce SAR

SA is a well-established regulator of SAR across multiple plant species, including Arabidopsis, tobacco, and cucumber (Zheng et al., 2015; Malamy et al., 1990; Métraux et al., 1990). Our findings, however, reveal that certain Brassicaceae species—*B. napus*, *B. rapa*, and *B. oleracea*—lack systemic SA accumulation (**Figure 5 and Supplementary Figure 3**), suggesting the absence of canonical systemic SA signaling in these species. To determine whether this deficiency compromises SAR, we conducted SAR assays, using the same conditions established for Arabidopsis (Zheng et al., 2015). The magnitude of SAR in these species was consistently lower than that observed in Arabidopsis Col-0 plants, highlighting the importance of systemic SA accumulation for robust SAR responses (**Figure 6**). Nonetheless, these species could still mount a modest SAR response in the absence of the CHE–*ICS1* regulatory module and systemic SA induction, pointing to the existence of alternative, SA-independent pathways of systemic immune signaling in these species. This highlights the evolutionary diversification of SAR regulatory mechanisms and underscores the need for further investigation into species-specific immune signaling networks.

**Figure 6.**
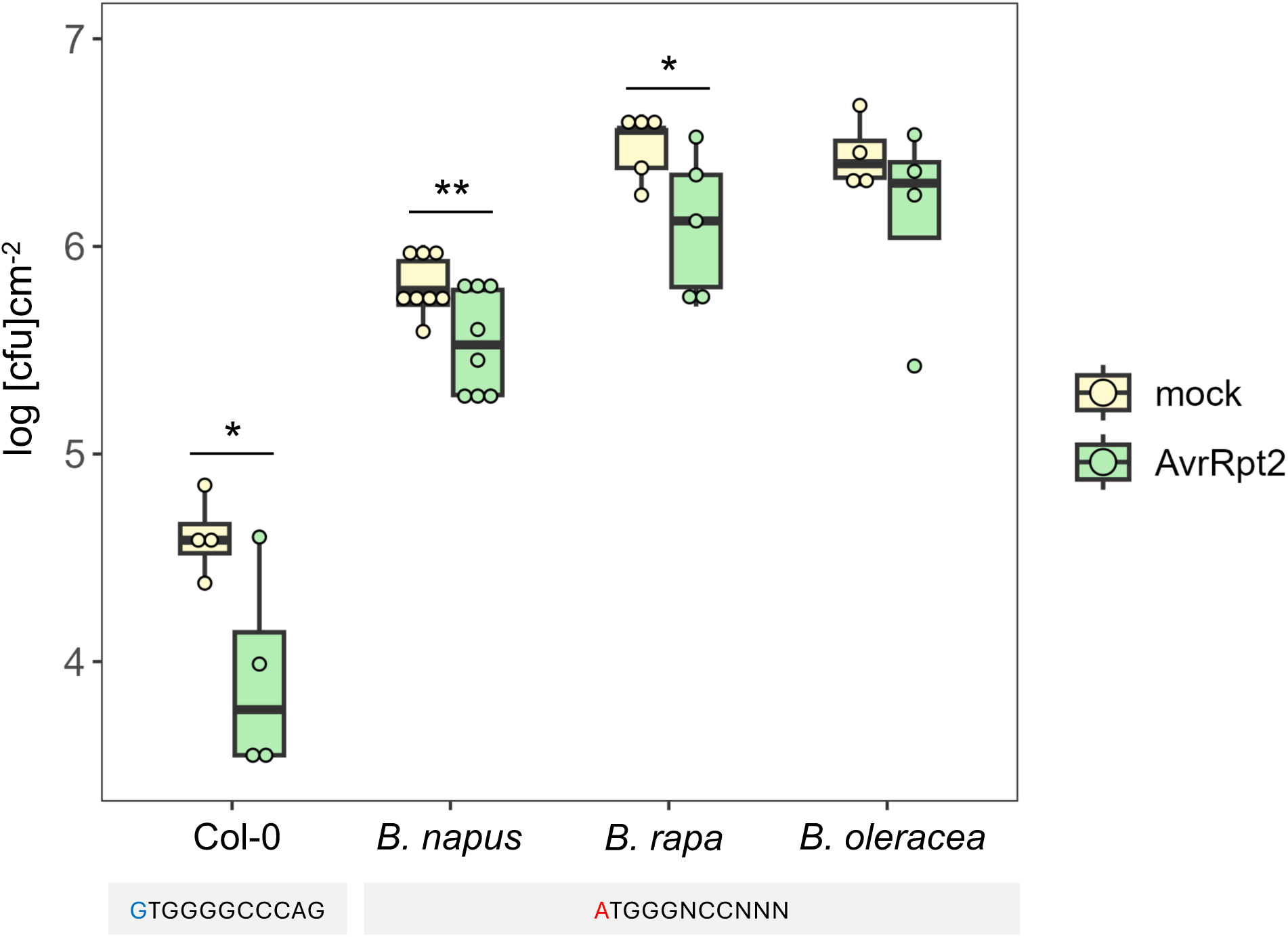
Modest SAR capacity is retained in species lacking the CHE–*ICS1* interaction. SAR was evaluated in Arabidopsis, *Brassica napus*, *B. oleracea*, and *B. rapa*. Plants were initially inoculated with MgCl_2_ (mock) or *Psm* ES4326/*AvrRpt2* (OD₆₀₀ = 0.02). Three days later, distal leaves were infected with *Psm* ES4326 (OD₆₀₀ = 0.001). Bacterial growth was measured three days after the second infection. All tested species showed reduced bacterial growth in systemic leaves following primary infection, indicating the establishment of SAR. However, the extent of the SAR was greater in Arabidopsis, which retains a canonical CHE–*ICS1* interaction (blue “G”), compared to Brassica species carrying a divergent TCP motif (red “A”), suggesting weaker SAR in the absence of systemic *ICS1* induction. Error bars represent the SE. Significant differences were calculated using two-tailed Student’s t-tests. \**p* < 0.1 and \*\**p* < 0.05.

### Introducing the CHE–*ICS1* module into *B. napus* enhances systemic SA accumulation

When exogenous SA was applied, *B. napus* exhibited significantly reduced bacterial growth (**Figure 7A**), consistent with previous studies demonstrating that elevated SA levels enhance disease resistance (Görlach et al., 1996; Nie, 2006). These results suggest that *B. napus* retains the ability to induce resistance in response to SA to establish SAR. To enable systemic SA induction and enhance SAR in this species we introduced a functional CHE-*ICS1* regulatory module. Specifically, the Arabidopsis *ICS1* coding sequence, driven by its native promoter (*pAtICS1*::AtICS1), was introduced into *B. napus* via *Agrobacterium*-mediated hypocotyl transformation (**Figure 7B**). Successful transformation was confirmed by PCR, which detected *AtICS1* gene fragments in the transgenic lines but not in wild-type *B. napus* (**Figure 7C**). Following local infection with *Psm* ES4326/*AvrRpt2*, transgenic *B. napus* exhibited upregulated *AtICS1* expression in systemic tissues at 6 hours post infection (hpi), whereas the expression of endogenous *BnICS1* remained unchanged (**Figure 7D** and **7E**). This transcriptional activation was followed by elevated SA levels at 12 hpi (**Figure 7F**). In contrast, this systemic response was absent in wild-type *B. napus* from as early as 6 hpi through 72 hpi (**Figure 5** and **7G**). Remarkably, the introduction of Arabidopsis *ICS1* carrying a canonical TCP binding motif successfully promoted early and robust systemic SA induction in *B. napus*. This result strongly suggests that native BnCHE proteins retain the ability to recognize the *AtICS1* promoter, as found in the Y1H assay (**Figure 3A**), and activate its expression in response to systemic signals. Thus, providing a single *cis*-regulatory element is sufficient to establish a functional CHE-*ICS1* regulatory module and enable systemic SA responses.

**Figure 7.**
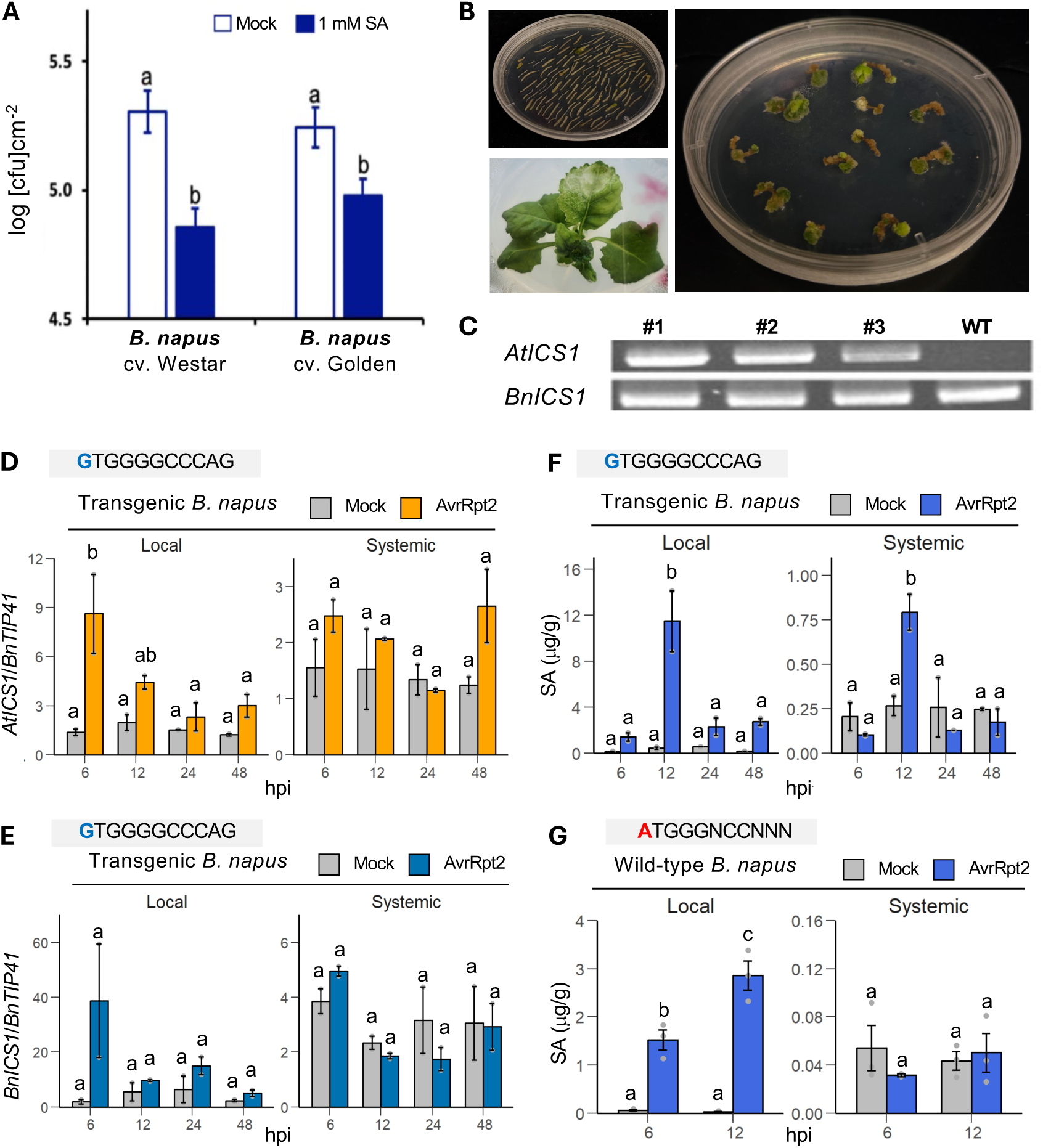
Introduction of the Arabidopsis CHE–*ICS1* module enables systemic SA accumulation in *B. napus.* **(A)** Exogenous SA treatment prior to infection with *Psm* ES4326 (OD₆₀₀ = 0.001) significantly reduced bacterial growth in both *B. napus* cvs. Westar and Golden. Plants were sprayed with 1 mM SA or water (control), and one day later, leaves were infected with *Psm*4326. Bacterial growth was measured 3 days post-infection (*n* = 8). Different letters indicate statistically significant differences (*p* < 0.05, one-way ANOVA followed by Tukey’s multiple comparisons test). **(B)** Arabidopsis *ICS1* under its native promoter (*pAtICS1*::*AtICS1*) was introduced into *B. napus* via *Agrobacterium-*mediated hypocotyl transformation. Representative images show explants, regenerating calli, and transgenic plantlets. **(C)** PCR confirmed the presence of *AtICS1* in three independent transgenic lines (#1–#3): wild-type *B. napus* (Bn) lacks *AtICS1*, while all lines, including wild-type, have *BnICS1* bands. **(D–E)** In transgenic *B. napus*, *AtICS1* expression was strongly induced in local tissue at all tested time points and in systemic tissue at 6 hours post-infection (hpi) with *Psm* ES4326/*AvrRpt2* infection, while *BnICS1* was induced only locally. **(F)** Transgenic *B. napus* exhibited robust local SA accumulation, with significant induction in systemic tissue particularly at 12 hpi. **(G)** In wild-type *B. napus* cv. Westar, while local SA accumulation was evident at both 6 and 12 hpi, systemic SA accumulation did not occur. Different letters indicate statistically significant differences (*p* < 0.05, one-way ANOVA followed by Tukey’s multiple comparisons test).

## Discussion

This study reveals how natural variation in a single *cis-*regulatory element can drive species-specific differences in systemic immune signaling by altering SA dynamics. The TCP-binding motif in the *ICS1* promoter, previously characterized in Arabidopsis, mediates the CHE-dependent activation of systemic SA biosynthesis. We identified sequence variations in this motif across plant species and demonstrated that this evolutionary divergence affects CHE binding, *ICS1* transcriptional activation, and systemic SA accumulation, leading to different levels of SAR across plant species. Consistent with the ICS1 pathway being the major SA biosynthetic route in the Brassicaceae, all tested Brassicaceae species exhibited strong *ICS1* induction and SA accumulation in locally infected tissue. However, only *C. sativa*, which retains a functional TCP-binding site, exhibited systemic *ICS1* expression and SA accumulation comparable to Arabidopsis. In contrast, *B. napus*, *B. rapa, and B. oleracea,* which all lack a canonical TCP-binding site, failed to induce *ICS1* and accumulate SA in systemic tissue. Since distinct transcriptional regulators mediate local and systemic *ICS1* expression—and since CHE–*ICS1* interaction was observed in *C. sativa* but not in *B. napus*— these findings provide a logical explanation for the distinct local and systemic responses in these species.

Our data also reinforce the central role of the ICS1 pathway in the Brassicaceae, as all tested species showed a clear correlation between *ICS1* expression and SA accumulation in both locally infected and systemic tissues. Especially in systemic tissues, species that exhibited *ICS1* induction also showed systemic SA accumulation, whereas those lacking *ICS1* expression did not accumulate SA. Although the PAL pathway is an alternative route for SA biosynthesis, the absence of *ICS1* induction and lack of SA accumulation in the systemic tissues of *B. napus, B. oleracea*, and *B. rapa* indicates that the PAL pathway does not contribute to systemic SA accumulation in these species. In non-Brassicaceae species, where the PAL pathway likely dominates in SA production (Farahani et al., 2016; Zhang et al., 2017; Zheng et al., 2019; Duba et al., 2019; Jia et al., 2023; Torrens-Spence et al., 2024; Hong et al., 2025), the systemic expressions of the pathway components, such as *PAL or AIM1* has not been well characterized, highlighting the need for further research. Therefore, to fully understand how a broad range of plants regulate their SA biosynthesis in local and systemic responses, it will be essential to investigate the systemic regulation of the PAL pathway through studies on *cis*-regulatory mechanisms and candidate transcription factors, as well as SA accumulation in non-Brassicaceae species. Unlike *ICS1*, which typically exists as a low-copy gene, *PAL* genes belong to large and diverse families with multiple paralogs in each species, making direct cross-species promoter comparisons more complex and challenging (Oyanagi et al., 2001; Zhan et al., 2022; Gao et al., 2023). This complexity highlights the need for tailored approaches when studying *PAL* regulation across different plant lineages, as well as the importance of investigating key genes encoding enzymes in this pathway once they are identified.

Studies from the past decade have reported pathogen-induced *ICS1* expression in several non-Brassicaceae crops, including pearl millet, rice, and barley (Siddaiah et al., 2017; Choi et al., 2015; Hao et al., 2018), suggesting that ICS1 may play a central role in SA biosynthesis in these species. However, more recent findings suggest that while the ICS1 pathway is the primary contributor to SA biosynthesis in the Brassicaceae, the PAL pathway may play a more dominant role in non-Brassicaceae species (Hong et al., 2025). This interpretation is supported by phylogenetic analyses showing the loss of some components downstream of ICS1 and the limited activity of associated enzymatic steps outside the Brassicaceae lineage (Hong et al., 2025). Although previous studies reported a correlation between *ICS1* expression and SA accumulation following pathogen attack (Siddaiah et al., 2017; Choi et al., 2015; Hao et al., 2018), it remains possible that PAL, rather than ICS1, was the primary contributor to SA accumulation in those cases. In this context, *ICS1* induction in non-Brassicaceae species may reflect an indirect response to stress or other signaling events rather than engagement of the ICS1-dependent pathway, a possibility that warrants further investigation. To date, direct genetic comparisons between the ICS1 and PAL pathways across species remain limited. A comprehensive analysis using robust comparative assays and functional genomics across diverse plant lineages will be critical to test whether recent models of ICS1 versus PAL usage are valid across species. Future studies quantifying pathway-specific gene expression, enzyme activity, and metabolite flux will be essential to elucidate the regulatory logic and evolutionary diversification of systemic SA biosynthesis.

Despite their lack of systemic SA induction, *B. napus* and other Brassica species still exhibited measurable but reduced SAR, indicating that the CHE–*ICS1* interaction in systemic SA biosynthesis is not strictly required for SAR. These results highlight the evolutionary flexibility of systemic immune networks and suggest the presence of alternative SAR signaling modules across species; thus, further investigation into the molecular basis of SA-independent SAR mechanisms is warranted. Indeed, in addition to SA, other signaling molecules—including Pip and NHP—are well-established mediators of SAR (Bernsdorff et al., 2016; Hartmann et al., 2018; Yildiz et al., 2021). Interestingly, recent work has shown that CHE regulates the expression of the Pip/NHP biosynthetic genes *ALD1*, *SARD4*, and *FMO1* (Cao et al., 2024), raising the possibility that SAR in some Brassica species may be mediated through Pip/NHP rather than SA and that CHE may contribute to SAR independently of *ICS1*. Future profiling of Pip and NHP across species will clarify whether these metabolites functionally compensate for impaired SA signaling. Moreover, amino-acid-derived signals such as serine have also been shown to promote immune priming (Lee et al., 2025). Expanding metabolomic analyses to include these molecules may uncover additional species-specific SAR signatures.

Our phylogenetic and sequence conservation analyses reveal that the TCP transcription factor family has diverged into 12 clades, with Clade A TCPs representing CHE orthologs in species ranging from lycophytes to angiosperms. However, within the Brassicaceae, Clade A TCPs have further diverged into CHE-like and TCP7-like subclades, suggesting the functional specialization of CHE in Brassicaceae species. This divergence raises the question of whether CHE’s role in activating systemic *ICS1* expression is an ancient trait of Clade A TCPs or a Brassicaceae-specific adaptation. In Arabidopsis, CHE’s ability to activate systemic *ICS1* expression depends on redox-sensitive sulfenylation at a conserved cysteine, which is highly conserved across Clade A TCPs (**Figure 1B**). Consistent with the importance of redox regulation, a CHE ortholog in *Nicotiana benthamiana* also undergoes sulfenylation in systemic tissues following local pathogen infection or exogenous H_2_O_2_ treatment (Cao et al., 2024), suggesting that the Clade A TCPs may act as redox-sensitive receivers of systemic signals across diverse plant lineages. Given that Arabidopsis CHE also regulates other components important for SAR, such as NHP biosynthetic genes (Cao et al., 2024), it will be important to investigate whether CHE homologs function as broader SAR regulators independent of the dominant SA biosynthetic route across species. Given that the sulfenylation of CHE enhances its binding to the *ICS1* promoter, and since the sulfenylation site is highly conserved across species (**Figure 1B**), it would be informative to test whether the species that exhibit systemic *ICS1* activation via CHE also utilize H₂O₂ as a mobile signal and whether this systemic regulation occurs through S-sulfenylation.

Beyond its role in systemic immune signaling, CHE also contributes to the circadian regulation of *ICS1* expression and SA biosynthesis in Arabidopsis, with SA levels peaking at dawn—a phase that coincides with increased pathogen risk—which may enhance immune preparedness (Wang et al., 2011; Goodspeed et al., 2012; Zheng et al., 2015; Karapetyan & Dong, 2018). Whether this circadian control is conserved in species that possess a functional CHE–*ICS1* module and missing in species that lack it remains an open question. Comparative time-course studies will be critical to determine whether the temporal regulation of SA represents a conserved immune strategy or an Arabidopsis-specific adaptation. Testing for changes in pathogen resistance in the morning versus nighttime across species would also be a valuable way to determine whether circadian timing can be exploited to enhance resistance across species.

Notably, introducing the Arabidopsis CHE–*ICS1* module into *B. napus* conferred systemic SA induction, firmly demonstrating *cis*-regulatory divergence in the *ICS1* promoter as the molecular basis for the deficiency of systemic SA accumulation. This successful gain of systemic transcriptional activation demonstrates the feasibility of engineering systemic immunity through targeted *cis*-regulatory modification. By enabling *de novo ICS1* synthesis in distal tissues, this strategy offers a promising path to enhance broad-spectrum disease resistance in Brassicaceae crops and potentially in other agriculturally important species. These findings present a proof of concept for transferring key transcriptional modules across species and highlight new opportunities for synthetic promoter engineering to rewire immune signaling networks. Importantly, these regulatory modules can be introduced without introducing full-length transgenes, such as the 2 kb Arabidopsis *ICS1* promoter. Instead, precise nucleotide-scale editing**—**such as an A-to-G substitution in the first position of the TCP binding site**—**could potentially establish functional promoter responsiveness, although further validation is required to confirm the effectiveness of this strategy. This targeted approach would not only reduce genomic disruption but also avoid the introduction of foreign DNA, offering advantages in public acceptance. Future efforts to optimize promoter strength and specificity could further enhance the robustness and precision of immune activation in crops.

## Methods

### Plant materials and growth conditions

All plants were grown in soil (Metro Mix 360) at 22 °C under a 12/12-h light/dark cycle with 55% humidity. *B. napus* cv. Westar and cv. Golden, *B. rapa* cv. K956, and *B. oleracea* cv. Crops-HH1 were obtained from U.S. National Plant Germplasm System. *C. sativa* cv. Celine was provided by Dr. Jeongim Kim at the University of Florida. *C. sativa* cv. Suneson was purchased from the Experimental Farm Network Seed Store (Philadelphia, PA, USA). *C. rubella* cv. Monte Gargano was purchased from Arabidopsis Biological Resource Center (CS22697). *C. annuum* cv. Hanyang was purchased from Holtgarden (Chino, California, USA).

### Identification and phylogenetic analysis of TCP proteins

Genome and gene annotation data for 56 species were obtained from Phytozome, NCBI, the Hornwort database, Ginkgo DB, and BnPIR (**Supplementary Table 1**). TCP-domain-containing sequences were identified using tBlastn searches (e-value cutoff: 1e^-10^), with the TCP domain sequences of Arabidopsis CHE (a Class I TCP) and TCP1 (a Class II TCP) used as query sequences. To construct the phylogenetic tree of the TCP transcription factor family, full-length TCP protein sequences were aligned using MAFFT v7.471 (Katoh and Standley, 2013). A maximum likelihood tree was inferred using IQ-TREE v3.0.1 (Wong et al., 2025) with JTT+F+R7 as the best fitting model. Branch support was assessed using both SH-aLRT and ultrafast bootstrap analyses with 1,000 replicates (Hoang et al., 2018). The resulting tree was visualized using iTOL (Letunic and Bork, 2024), and TCP proteins were classified into 12 major clades based on topological relationships. Conservation within the Clade A TCPs was assessed by calculating per-residue conservation scores using JalView v2.11.4.1 (Waterhouse et al., 2009), following a multiple sequence alignment with MAFFT v7.471.

### Identification of ICS1 and phylogenetic analysis of ICS1 promoters

Homologs of Arabidopsis *ICS1* were identified via tBLASTn searches and manually curated to confirm the presence of a chloroplast transit peptide using Target P 2.0 (Almagro Armenteros et al., 2019), as ICS1 localizes to the chloroplast to function properly (Garcion et al., 2008). The phylogenetic analysis of *ICS1* promoter regions was performed using approximately 2 kb of the genomic sequences upstream of the *ICS1* coding region. Sequences were aligned using MAFFT v7.471, and a maximum likelihood tree was generated using IQ-TREE v3.0.1 with TVM+F+R2 as the best fitting model.

### Identification of TCP-binding sites within *ICS1* promoters

Promoter regions corresponding to 2 kb upstream of the *ICS1* coding sequences were scanned for TCP-binding motifs using the FIMO tool in MEME Suite 5. 5. 7 (Charles et al, 2011), with the canonical CHE binding motif sequence obtained from the Plant Transcription Factor Database (Tian et al., 2020). Identified TCP-binding motif were categorized based on sequence variation and mapped onto the phylogenetic tree of *ICS1* promoters to assess lineage-specific divergence in the *cis*-regulatory elements (**Figure 2**).

### Cloning of plasmid constructs

For the Y1H assays, *ICS1* promoters from *A. thaliana* Col-0, *B. napus* cv. Westar, *C. sativa* cv. Suneson, and *C. annuum* cv. Hanyang were amplified via PCR using the primers listed in **Supplementary Table 2**. The promoter fragments were assembled into SalI/SacI-digested pABAi using NEB HiFi DNA Assembly Master Mix. *CHE* coding sequences from the corresponding species were amplified via PCR and cloned into pGADT7 digested with EcoRI and BamHI to generate fusions with the Gal4 activation domain. To clone all *BnCHE* homologs, including *BnCHE_A*, *BnCHE_B*, *BnCHE_C*, *BnCHE_D*, *BnCHE_E*, and *BnCHE_F*, UTR-specific primers were designed due to their highly conserved coding sequences. DH5-α competent *Escherichia coli* was used as a host to transform the *pABAi-ICS1* and *pGADT7-CHE* plasmid constructs. The transformants were selected by plating the bacterial suspension on LB media containing 50 µg/mL carbenicillin and incubating overnight at 37°C. Single colonies were picked and subjected to PCR to identify transformed colonies. For the protoplast-based GUS reporter assay, the same set of *ICS1* promoters used in the Y1H assay and the Arabidopsis CHE (AtCHE) coding sequence were amplified by PCR using the corresponding primers (**Supplementary Table 2**). To generate the reporter constructs, the *ICS1* promoters were cloned upstream of the GUS reporter gene in the pMIN35S vector. For the effector construct, the AtCHE coding sequence was fused to the sCFP tag and inserted into the p326 vector under the control of the CaMV 35S promoter. The empty vector control was generated by inserting an sCFP tag into the p326 vector without CHE.

### Plant transformation

To generate Arabidopsis transgenic lines carrying *ICS1* genes driven by various promoters, including the native promoter from Arabidopsis, G-to-A mutated Arabidopsis, *C. sativa*, and *B. napus*, genomic *ICS1* fragments, including the entire coding region and approximately 2 kb of the upstream sequence, were PCR-amplified from Arabidopsis Col-0, *C. sativa* cv. Suneson, and *B. napus* cv. Westar. The amplified fragments were cloned into the binary vector pJHA212G-RBCSt. The resulting constructs were introduced into *Agrobacterium tumefaciens* strain GV3101 to transform the Arabidopsis *sid-2* mutant via floral dipping method (Clough et al., 1998). More than 15 T1 lines were generated for each transgenic line, and two homozygous lines were selected for each construct. To generate transgenic *B. napus* lines expressing Arabidopsis *ICS1* under the control of the Arabidopsis *ICS1* promoter, the same genomic *ICS1* construct used for the Arabidopsis transgenic lines, including the entire coding region and approximately 2 kb of the upstream sequence from Arabidopsis Col-0, were PCR amplified and used. *B. napus* cv. Westar was transformed via Agrobacterium-mediated hypocotyl transformation, as previously described (Dai et al., 2020).

### Yeast one-hybrid assay

The *pABAi-ICS1* plasmid construct was digested with the BbsI restriction enzyme and integrated into the genome of the Y1HGold yeast strain. The transformed yeast was grown on SD/-uracil medium, and a single colony was selected for subsequent transformation with the *CHE* expression vector. The resulting yeast strain containing both the *ICS1* promoter and *CHE* was grown on SD/-leucine medium, and a single colony was selected for the Y1H assay to test CHE binding to the *ICS1* promoter. To assess the interaction, the colony was cultured overnight in 5 mL SD/-leucine medium. The next day, yeast cells were plated on SD/-leucine medium (control) and SD/-leucine supplemented with Aureobasidin A (AbA) (selective medium) in parallel.

### Protoplast transient expression and GUS reporter assay

Four-week-old Arabidopsis Col-0 and *che-2* were infiltrated with *Psm* ES4326/*AvrRpt2* (OD_600nm_=0.02) or mock-treated with 10 mM MgCl_2_. Two days after infection, protoplasts were isolated from systemic uninfected tissues using an enzyme solution (1.5% cellulase, 0.4% macerozyme, 0.4 M mannitol, 10 mM CaCl_2_, 20 mm KCl, 20 mM 2-(*N*-morpholine)-ethanesulfonic acid (MES; pH 5.7), and 0.1% bovine serum albumin). The protoplasts were then filtered through a 40-μm stainless mesh and collected in W5 solution (154 mM NaCl, 125 mM CaCl_2_, 5 mM KCl, 2 mM MES and 5 mM glucose; pH 5.7). Protoplasts were then resuspended in MMg solution (0.4 M D-mannitol, 15 mM MgCl_2_, and 4 mM; pH 5.7). For transfection, a 1:1 volume of 40 % polyethylene glycol (PEG) solution containing 0.2 M D-mannitol and 100 mM CaCl_2_ was added to the protoplast suspension with reporter and effector plasmids and incubated for 15 min. A CaMV 35S promoter-driven luciferase construct was also co-transfected as an internal control. The protoplasts were resuspended in W5 solution and incubated overnight at dark. Transformed protoplasts were lysed in Cell Culture Lysis Reagent (Promega) for downstream assays. For GUS activity measurement, 50 μl of lysate was incubated with 142 μl of GUS extraction buffer (1 M NaH_2_PO_4_, 0.5 M EDTA, 0.1 % Triton X-100, and 0.5 % beta-mercaptoethanol) and 8 μl of 25 mM 4-methylumbelliferyl-beta-D-glucuronide at 37 °C for 1 h. After incubation, 40 μl of the reaction mixture was transferred to a black 96-well plate, and 200 μl of 0.2 M Na_2_CO_3_ was added to stop the reaction. GUS fluorescence was measured using a Tecan microplate reader (Bio-Tek) with excitation at 360 nm and emission at 465 nm. For luciferase activity, 40 ul of Luciferase Assay Buffer (Promega) was added to 40 ul of the cell lysate in a white 96 well plate, and luminescence was measured using the same microplate reader.

### Chromatin immunoprecipitation assay

The ChIP assay was performed as previously described (Lee et al., 2021) with minor modifications. The relative abundance of specific DNA fragments before and after the immunoprecipitation were quantified by qPCR using primer listed in **Supplementary Table 2**. Percent input values were calculated as the ratio of DNA abundance in the immunoprecipitated sample to that in the input (pre-immunoprecipitation) sample. The percent input values for each target region (e.g., a TCP-binding site in the *ICS1* promoter) were normalized to control genomic regions: *PP2A and EIF4a* for Arabidopsis, *TIP41* and GADPH for *B. napus*, *TIP41* for *B. rapa*, *UBQ* for *C. rubella*, and *ACT100* for *C. annuum*.

### Site-directed mutagenesis

Site-directed mutagenesis was performed using overlap extension PCR to introduce a G-to-A substitution at the first nucleotide of the TCP-binding site in *AtICS1* and *CsICS1_A.* Similarly, glycine-to-lysine substitutions were introduced at positions G68 in AtCHE and G70 in BnCHE by changing GGT to AAG and GGC to AAG, respectively. For each mutation, two pairs of primers were designed such that the mutation was located at the center of the reverse primer of pair 1 and of the forward primer of pair 2. These primer pairs generated two overlapping PCR fragments, which were assembled in a second round of PCR using the forward primer of pair 1 and the reverse primer of pair 2 to produce a single fragment containing the desired substitution. All mutations were verified by Sanger sequencing. The resulting mutated *ICS1* and *CHE* fragments were then cloned into the pABAi and pGADT7 vectors, respectively, for Y1H assays. The mutated *AtICS1* constructs were also cloned into the pMIN35S and pJHA212G-RBCSt vectors for the protoplast-based GUS reporter assay and plant transformation, respectively.

### Quantitative real-time reverse transcription-polymerase chain reaction (qRT-PCR)

Total RNA was isolated using a Quick-RNA Miniprep Kit (Zymo Research) following the manufacturer’s protocol. First-strand cDNA was synthesized from 1 µg of total RNA using a LunaScript™ RT SuperMix Kit (New England Biolabs). Quantitative PCR was carried out using a LightCycler® 480 system (Roche) with Luna® Universal qPCR Master Mix and primers listed in **Supplementary Table 2**. For species with multiple gene homologs, primers were designed to target conserved regions to ensure detection across orthologs. Transcript levels were normalized to *TIP41* as a validated reference gene (Dixit et al., 2019).

### Salicylic acid measurement and analysis

SA levels and the SA storage form SA glucoside (SAG) were analyzed using high-performance liquid chromatography (HPLC) as previously described (Zheng et al., 2015). Briefly, 50–100 mg of leaf tissue was ground and extracted with 90% methanol. The mixture was centrifuged for 10 minutes, and the supernatant (first extract) was transferred to a fresh tube. The remaining pellet was re-extracted with 100% methanol and centrifuged again for 10 minutes, and the resulting supernatant was combined with the first extract. The combined extract was centrifuged once more, and the final supernatant was vacuum-dried at 45 °C for 3 hours. The dried residue was resuspended in 5% (v/v) trichloroacetic acid. To extract free SA, samples were subjected to two rounds of extraction with ethyl acetate–cyclopentane (1:1, v/v), followed by drying at 45 °C for 40–45 minutes and resuspension in HPLC elution buffer [10% (v/v) methanol in 0.2 M sodium acetate buffer]. SA was analyzed using an Agilent 1260 HPLC system equipped with an XTerra MS C18 column (3.5 µm, 3.0 × 100 mm). Peak areas were compared against SA standard curves to determine absolute concentrations.

### SAR test

For the SAR test, 3-week-old plants were infiltrated with *Psm*ES4326/*AvrRpt2* (OD_600nm_ = 0.02) to induce systemic immunity. 10 mM MgCl₂ was infiltrated as a control. After 3 days, systemic leaves from previously infected or mock-treated plants were infiltrated with the virulent bacterial pathogen *Psm*ES4326 (OD_600nm_ = 0.001). Leaf discs from systemic leaves were then collected at 3 days post infiltration. Leaf discs were ground and resuspended in 10 mM MgCl_2_ and plated on King’s B (KB) medium with serial dilutions. Bacterial colonies were scored after plating and incubating at 30°C for two days.

### Statistical analysis

For statistical analyses, two-tailed Student’s t-tests and one-way ANOVA followed by Tukey’s multiple comparisons test were performed to determine significant differences. Details of each analysis are given in the figure legends.

## Acknowledgments

We thank Dr. Xinnian Dong for providing the polyclonal AtCHE antibody. Special thanks to former lab members Dr. Saborni Maiti and Samuel Marston for their initial contributions to salicylic acid quantification and *Agrobacterium*-mediated transformation of *Brassica napus*, respectively. We also thank members of the HYoo lab for their valuable feedback throughout this project. This work was supported by start-up funds from the University of Utah and Oklahoma State University.

## Author Contributions

J.H., R.A., C.Y., and H.Y. designed the research. J.H. performed the evolutionary analysis and the computational analyses. R.A. generated constructs for the Y1H assays, including the mutated *ICS1* promoters and CHEs. R.A. and C.Y. performed the Y1H assays. J.H., and C.Y. carried out the qRT-PCR analyses. J.H., H.D.R., M.L.D.F., H.W., and A.M.C. conducted the salicylic acid measurements. J.H., H.D.R., M.L.D.F., H.A., and A.N. performed the SAR test. R.A. conducted the *Agrobacterium*-mediated transformation using the floral dip method to generate transgenic Arabidopsis plants, and M.L.D.F. conducted the *Agrobacterium*-mediated transformation of *Brassica napus* hypocotyls. J.H. performed all other experiments, including ChIP-qPCR, protoplast-based transient expression assays to evaluate CHE–*ICS1* interactions *in vivo*, western blot, and experiments involving *B. napus* transgenic plants. J.H. and H.Y. wrote the manuscript with input from all authors.

---

**Supplementary Table 1.** Genome information of 53 species and the number of identified TCP proteins

**Supplementary Table 2.** Primers used in this study

## Notes

### Competing Interest Statement

The authors have declared no competing interest.

