## Supplementary Tables for "*Cis*-regulatory variation in *ISOCHORISMATE SYNTHASE 1* modulates systemic salicylic acid biosynthesis and systemic acquired resistance in plants"

**Supplementary Table 1.** Genome information of 53 species and the number of identified TCP proteins

| **Classification** | **Species** | **Abbr.** | **Source** | **Version** | **TCP** |
| --- | --- | --- | --- | --- | --- |
| Rhodophyta | *Porphyra umbilicalis* |  | Phytozome | v1.5 | **0** |
| Chlorophyte | *Botryococcus braunii* |  | Phytozome | v2.1 | **0** |
|  | *Chlamydomonas reinhardtii* |  | Phytozome | v5.6 | **0** |
|  | *Coccomyxa subellipsoidea* |  | Phytozome | v2.0 | **0** |
|  | *Micromonas pusilla* CCMP1545 |  | Phytozome | v3.0 | **0** |
|  | *Micromonas* sp RCC299 |  | Phytozome | v3.0 | **0** |
|  | *Ostreococcus lucimarinus* |  | Phytozome | v2.0 | **0** |
|  | *Chlamydomonas reinhardtii* |  | Phytozome | v5.6 | **0** |
|  | *Volvox carteri* |  | Phytozome | v2.1 | **0** |
|  | *Dunaliella salina* |  | Phytozome | v1.0 | **0** |
|  | *Chromochloris zofingiensis* |  | Phytozome | v5.2.3.2 | **0** |
| Streptophyte Algae | *Mesostigma viride* |  | NCBI | GCA_009746045.1 | **0** |
|  | *Klebsormidium nitens* |  | NCBI | GCA_000708835.1 | **0** |
|  | *Chara braunii* |  | NCBI | GCA_003427395.1 | **2** |
| Hornworts | *Anthoceros agrestis* |  | Hornwort database |  | **2** |
| Liverworts | *Marchantia polymorpha* |  | Phytozome | v3.1 | **2** |
| Moss | *Sphagnum fallax* |  | Phytozome | v1.1 | **5** |
|  | *Physcomitrella patens* |  | Phytozome | v3.3 | **6** |
|  | *Ceratodon purpureus* |  | Phytozome | R40 v1.1 | **1** |
| Lycophyte | *Selaginella moellendorffii* | Selmo | NCBI | GCF_000143415.4 | **12** |
| Fern | *Ceratopteris richardii* | Cerric | Phytozome | v2.1 | **13** |
| Gymnosperm | *Ginkgo biloba* |  | GinkgoDB |  | **13** |
|  | *Thuja plicata* |  | Phytozome | v3.1 | **17** |
| basal angiosperm | *Amborella trichopoda* | Ambtr | NCBI | GCF_000471905.2 | **10** |
|  | *Nymphaea colorata* | Nymco | Phytozome | V1.2 | **16** |
|  | *Quercus lobata* | Quelo | NCBI | GCF_001633185.2 | **21** |
|  | *Liriodendron tulipifera* | Lirtu | Phytozome | V1.1 | **16** |
| monocot | *Brachypodium distachyon* | Bradi | Phytozome | v3.2 | **20** |
|  | *Zea mays* | Zeama | Phytozome | BB4 v1.2 | **32** |
|  | *Setaria viridis* | Setvi | Phytozome | v2.1 | **22** |
|  | *Sorghum bicolor* | Sorbi | Phytozome | v3.1.1 | **19** |
|  | *Oryza sativa* | Orysa | Phytozome | v7.0 | **20** |
|  | *Spirodela polyrhiza* | Spipo | Phytozome | v2.0 | **16** |
|  | *Zostera marina* | Zosma | Phytozome | v3.1 | **14** |
| Dicot | *Vitis vinifera* | Vitvi | Phytozome | v2.1 | **20** |
|  | *Cucumis sativus* | Cucsa | NCBI | GCF_000004075.3 | **27** |
|  | *Glycine max* | Glyma | Phytozome | Wm82.a4.v1 | **54** |
|  | *Medicago truncatula* | Medtr | Phytozome | Mt4.0v1 | **21** |
|  | *Populus trichocarpa* | Poptr | Phytozome | v3.0 | **34** |
|  | *Carica papaya* | Carpa | Phytozome | ASGPBv0.4 | **21** |
|  | *Theobroma cacao* | Theca | Phytozome | v1.1 | **21** |
|  | *Arabidopsis thaliana* | Arath | Phytozome | TAIR10 | **24** |
|  | *Arabidospis lyrata* | Araly | Phytozome | v2.1 | **25** |
|  | *Camelina sativa* | Camsa | NCBI | GCF_000633955.1 | **68** |
|  | *Capsella rubella* | Caprub | Phytozome | v1.1 | **23** |
|  | *Eutrema salsugineum* | Eutsa | NCBI | GCF_000478725.1 | **24** |
|  | *Brassica oleracea* | Braole | NCBI | GCF_000695525.1 | **41** |
|  | *Brassica rapa* | Brara | NCBI | GCF_000309985.2 | **41** |
|  | *Brassica napus* | Branap | BnPIR | Westar | **82** |
|  | *Solanum lycopersium* | Solly | Phytozome | ITAG2.4 | **36** |
|  | *Solanum tuberosom* | Soltu | Phytozome | v6.1 | **32** |
|  | *Nicotiana tabacum* | Nicta | NCBI | GCF_00071513 | **66** |
|  | *Solanum pennelli* |  | NCBI | GCF_001406875.1 |  |
|  | *Capsium annuum* | Capan | NCBI | GCF_002878395.1 | **29** |

**Supplementary Table 2.** Primer list

| **Yeast one-hybrid assay** | |
| --- | --- |
| **Name** | **Sequence** |
| AtICS1pro-F | AAGCTTGAATTCGAGCTAAAAATAAAAATTATATAGATATA |
| AtICS1pro-R | AGCACATGCCTCGAGGTTTGGGTTTAGAGTTGAG |
| CsICS1Apro-F | GAAAAGCTTGAATTCGAGCTACTCAAAGGAGTTGACTCC |
| CsICS1Apro-R | AGCACATGCCTCGAGGTGGAGAAATGAATTCTTTGGTTG |
| BnaICS1AEpro-F | GAAAAGCTTGAATTCGAGCTAGTCTCCTCTCACTCTTG |
| BnaICS1AEpro-R | AGCACATGCCTCGAGGTGGAGATTGCTTAGAGAACG |
| BnaICS1Bpro-F | AAAGCTTGAATTCGAGCTAAACTGGTGACCCGGTTC |
| BnaICS1Bpro-R | AGCACATGCCTCGAGGTGGAGATAGCTTAGAGAAC |
| BnaICS1CDpro-F | GAAAAGCTTGAATTCGAGCTACAATCCAGAACTGAACC |
| BnaICS1CDpro-R | AGCACATGCCTCGAGGTGGAGAACGAAAGAAACAG |
| CaICS1pro-F | GAAAAGCTTGAATTCGAGCTATGGTCATCTGATTTGGACGA |
| CaICS1pro-R | AGCACATGCCTCGAGGTTAAGATGGGGGACTGGACTG |
| AD-AtCHE-F | GCCATGGAGGCCAGTGCTATGGCCGACAACGACGGA |
| AD-AtCHE-R | GCCATGGAGGCCAGTGCTATGTCGACGTCTGCAGAAGG |
| AD-CsCHE12-F | GCCATGGAGGCCAGTGCTATGGCGAACAACGACGGA |
| AD-CsCHE12-R | CAGCTCGAGCTCGATGTCAACGTGGGTCGTGGTC |
| BnaCHE-UTR-AB-F | AACTGAACTCCCAAATCCAC |
| BnaCHE-UTR-AB-R | CTATACATACCCAAATGGAACC |
| BnaCHE-UTR-C-F | TCTTCTCGAGAGAGAGAGTG |
| BnaCHE-UTR-C-R | CTCAAACTAAAGTAGTCTCTG |
| BnaCHE-UTR-DE-F | TTACCTCTCAAATTCTCTCTTCC |
| BnaCHE-UTR-D-R | cacAaattaagtactcgttaaac |
| BnaCHE-UTR-E-R | CAGATTAAGTACTCGTTAAACC |
| BnaCHE-UTR--F-F | CAACCTTCTTCTTCTCGAGAG |
| BnaCHE-UTR-F-R | CTCGAACAGTCTCTGACC |
| AD-BnaCHE-AB-F | GCCATGGAGGCCAGTGCTATGTCGAACGACGACGG |
| AD-BnaCHE-A-R | CAGCTCGAGCTCGATGTCAACGAGAGTTATCCTCC |
| AD-BnaCHE-B-R | CAGCTCGAGCTCGATGCTAGCTGCCGAAGCCTCACC |
| AD-BnaCHE-CF-F | GCCATGGAGGCCAGTGCTATGGAGAACAACGACGGA |
| AD-BnaCHE-C-R | CAGCTCGAGCTCGATGTCAACGTGAGTCATCCTC |
| AD-BnaCHE-DE-F | GCCATGGAGGCCAGTGCTATGACGAGCAACGACGG |
| AD-BnaCHE-DE-R | CAGCTCGAGCTCGATGTTAACGTGGGTTATCCTCT |
| AD-BnaCHE-F-R | CAGCTCGAGCTCGATGTCAGCGTGAGTCATCCTC |
| AD-CaCHE-F | GCCATGGAGGCCAGTGCTATGTCGACGTCTGCAGAAGG |
| AD-CaCHE-R | CAGCTCGAGCTCGATGTCAGCGCCCGTCATCGTC |
| AD-BrCHE_F | GCCATGGAGGCCAGTGCTATGACGAGCAACGACGG |
| AD-BrCHE_R | AGCTCGAGCTCGATGTTAACGTGGGTTATCCTCTTCCCTC |
| AD-BoCHE1_F | GCCATGGAGGCCAGTGCTATGACGAGCAACGACGGA |
| AD-BoCHE1_R | AGCTCGAGCTCGATGTTAACGTGGGTTATCCTCTTCCCTC |
| AD-BoCHE2_F | GCCATGGAGGCCAGTGCTATGACGAGCAACGACGGA |
| AD-BoCHE2_R | AGCTCGAGCTCGATGTCAACGTGAGTCATCCTCCTCC |
| AD-BoCHE3_F | GCCATGGAGGCCAGTGCTATGGAGAACAACGACGGAGC |
| AD-BoCHE3_R | AGCTCGAGCTCGATGTCAACGTGAGTCATCCTCTTCCC |
| **Site-directed mutagenesis** | |
| **Name** | **Sequence** |
| AtICS1_GtoA_F | TTCAATTTTAaTGGGCCCCTG |
| AtICS1_GtoA_R | GATTTTCATTTCATTTTCACACAAAATTTC |
| CsICS1_GtoA_F | TCCAAAATTAaTGGGGCCCAG |
| CsICS1_GtoA_R | GATTCAGTTCACCTTTCACATTTTTAAATTTC |
| AtCHE-GtoK-F | CAAGTCCGATGGTCAAACCATA |
| AtCHE-GtoK-R | CTATGGTTTGACCATCGGACTTG |
| BnCHE_A (Glycine to lysine)_forward primer | CAAATCCGACAAGCAGACAATC |
| BnCHE_A (Glycine to lysine)_Reverse primer | GATTGTCTGCTTGTCGGATTT |
| **Protoplast based GUS reporter assay** | |
| **Name** | **Sequence** |
| AtCHE-p326-F | agaacacgggggactATGGCCGACAACGACGG |
| AtCHE-p326-R | ttccaccgcctcCaccACGTGGTTCGTGGTCG |
| AtICS1pro-pMIN35-F | AGCTCGGTACCCGGGATTCGTAGCATCCACAACACAC |
| AtICS1pro-pMIN35-R | ccttatatagaggaagggtcttaTGCAGAAATTCGTAAAGTGTTTCTTGA |
| CsICS1Apro-pMIN35-F | AGCTCGGTACCCGGGACTCAAAGGAGTTGACTCCTCAA |
| CsICS1Apro-pMIN35-R | ccttatatagaggaagggtcttaTGGAGAAATGAATTCTTTGGTTGATTTTTT |
| BnICS1Apro-pMIN35-F | AGCTCGGTACCCGGGAGTCTCCTCTCACTCTTGACG |
| BnICS1Apro-pMIN35-R | ccttatatagaggaagggtcttaTGGAGATTGCTTAGAGAACGAACA |
| **ChIP assay** | |
| **Name** | **Sequence** |
| AtICS1p-TCP-F | TGCACGACTAACTTTAGAAAAATGT |
| AtICS1p-TCP-R | AGGGGACTGATGTAGCAGGGGC |
| AtPP2A-gDNA-F | TATCGGATGACGATTCTTCGTGCAG |
| AtPP2A-gDNA-R | TTAGATGCAGTTATTACCGCAGG |
| AtEIF4a-gDNA-qF | GGTGTGTGTTAGTATGCAATTGC |
| AtEIF4a-gDNA-qR | CCCACTTTCCTCTTCCTTTTCG |
| CsICS1pAll-TCP-F | AAATGTGAAAGGTGAACTGAATC |
| CsICS1pAll-TCP-R | AGAGATGGGTTATACAATGGATTTG |
| CsTIP41-gDNA-F | GACATCTCACTTACCTGAAATGG |
| CsTIP41-gDNA-R | TGCAGCAGGAACTTCAACAG |
| CsEIF4A3-gDNA-F | CCAGCTTCTTCCTTCCAAGG |
| CsEIF4A3-gDNA-R | GAATCCTGACTGGTTTGTTCATG |
| CrICS1p-TCP-F | AGAAGTACGTATAGATAGTTTACAG |
| CrICS1p-TCP-R | CTAAAGATGAGTATACATTGTGTGC |
| CrUBQ-gDNA-F | tggcaagaccattactcttgag |
| CrUBQ-gDNA-R | CTGTTTACCGGCAAAGATCAATC |
| BnICS1pAE-TCP-F | AATTCTTCAATTTTTAAAATGGGGC |
| BnICS1pAE-TCP-R | ATGGATTATGGATTCGTCCCTC |
| BnICS1pB-TCP-F | GGTAACGAACATTTCTGAGCAG |
| BnICS1pB-TCP-R | GGCACGTCTTAAATAAGATGGG |
| BnICS1pCD-TCP-F | GTGGGACGGTGGAATGAATC |
| BnICS1pCD-TCP-R | GTAGTAAGAGACTAGAGATTTGAGC |
| BnTIP41-gDNA-F | AGGAGGTAAGAGTTTGTGATGAG |
| BnTIP41-gDNA-R | GTAATCATAATCCAGTATCACCTGC |
| BnGADPH-gDNA-F | TCTGAGATTTGAATCTTCGTGTAGG |
| Bn-GADPH-gDNA-R | GACGAGCTCAACATCGTTCC |
| BrICS1pA-TCP-F | agtgggaaggtggaatgaatc |
| BrICS1pA-TCP-R | GTCTTAAATAGACGTGACGGGG |
| BrICS1pB-TCP-F | AAGTATGATCCAATTTCAAGTGAAG |
| BrICS1pB-TCP-R | TTCAATAAATAGGGGACGGGTG |
| BrTIP41-gDNA-F | gatcgcaaatgtgaaaccgac |
| BrTIP41-gDNA-R | TAGAATCTCCTTATCGACCTCCG |
| CaICS1p-TCP1-F | GGTAGTTGGTGCCTGTGAATAG |
| CaICS1p-TCP1-R | TTTGTCATTGATAACCTTCGTTACC |
| CaICS1p-TCP2-F | GGGTTGCAACTTGTAATGACG |
| CaICS1p-TCP2-R | CTAGTTACTCTTGGAGCTACGAG |
| CaACT100-gDNA-F | cgtgtgtttatttgcaggatgt |
| CaACT100-gDNA-R | AAGTAGGGAAGCAAGAATTTCAAC |
| **qPCR** | |
| **Name** | **Sequence** |
| CsBn_TIP41qF | GGTTGAGAGAGACGAGAATGC |
| CsBn_TIP41qR | ACTGGATACCCTTTCGCAGA |
| CsICS1-qF | GCAGTGTTCTTCCGTGACCTTG |
| CsICS1-qR | AGGCACCAAAAGGTTCCCATTC |
| BnaICS1all-qF | CAGTTACAGAGTGAAGGGC |
| BnaICS1all-qR | AAGTCTCTCAGGCGTGTT |
| BrTIP41-qF | GTAGCGGAGTTGTTGAGAAAGA |
| BrTIP41-qR | CTCTTCACAGTTCTCCCACTTC |
| BrICS1all-qF | TGCACAGTTACAGAGTGAAGG |
| BrICS1all-qR | TGCAGACACCTAACTGATTCC |
| BoTIP41-qF | CCATGACTGGGAGATCGAAAC |
| BoTIP41-qR | GCCAAACACCATTTCAGGTAAG |
| BoICS1all-qF | GGAGATCCATCCGAAGGTTTC |
| BoICS1all-qR | TCCACAGAGATCTTGCCGTT |
| AtICS1-qF | GGCAGGGAGACTTACG |
| AtICS1-qR | AGGTCCCGCATACATT |
| **Plant transformation** | |
| **Name** | **Sequence** |
| AtICS1pro-1.748kb-comp-F | GAATTCGAGCTCGGTACATTCGTAGCATCCACAACACAC |
| AtICS1pro-1.748kb-comp-R | TGAAGCCATTGCAGAAATTCGT |
| AtICS1-CDS-comp-F | ATTTCTGCAATGGCTTCACTTCAA |
| AtICS1-CDS-comp-R | cgtatgggtaTTGTGAGAACCCCTTATCCCC |
| CsICS1pro-1.538kb-comp-F | GAATTCGAGCTCGGTACACTCAAAGGAGTTGACTCC |
| CsICS1pro-1.538kb-comp-R | GACGCCATTGGAGAAATGAATTCTTTGGTTG |
| CsICS1-CDS-comp-F | tttCTCCAATGGCGTCGCTTC |
| CsICS1-CDS-comp-R | ggaacatcgtatgggtaATTAATCGCATGCAGAGC |
| BnICS1pro-1.526kb-comp-F | GAATTCGAGCTCGGTACATTCGTAGCATCCACAACACAC |
| BnICS1pro-1.526kb-comp-R | TGAGGCCATTGGAGATTGCTTAGAGAACG |
| BnICS1-CDS-F | AATCTCCAATGGCCTCACTTCATC |
| BnICS1-CDS-R | ggaacatcgtatgggtaATTAATTGGCTGCAGAGTTGTTG |
