## Supplementary Figures for "*Cis*-regulatory variation in *ISOCHORISMATE SYNTHASE 1* modulates systemic salicylic acid biosynthesis and systemic acquired resistance in plants"

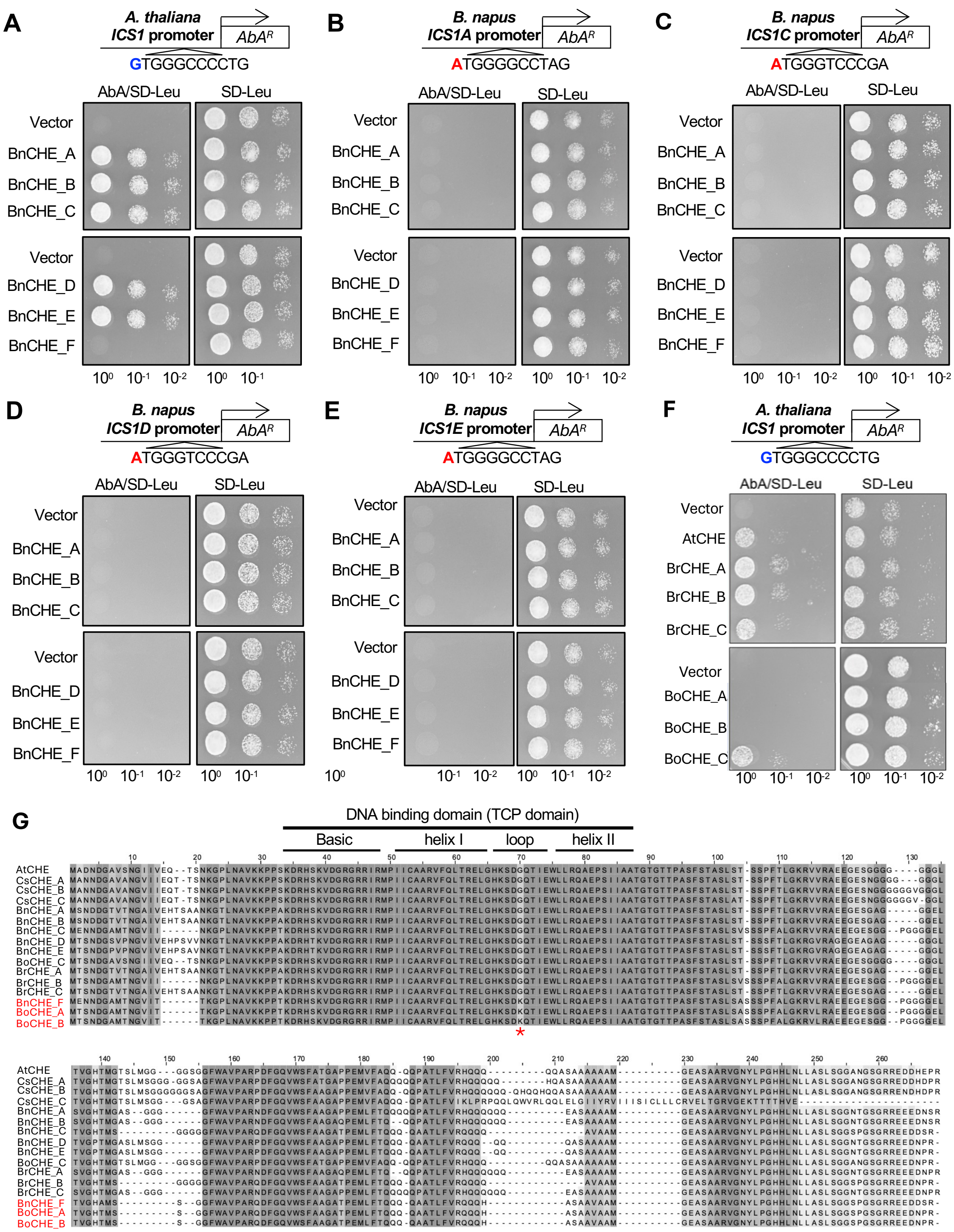


**Supplementary Figure 1**. **Analysis of CHE binding to *ICS1* promoters using CHE orthologs from *B. napus*, *B. rapa*, and *B. oleracea*. (A-E)** Yeast one-hybrid assays testing the interactions between individual *B. napus* CHE orthologs and *ICS1* promoters from Arabidopsis (A) and *B. napus* (B-E): *BnICS1_A* (B), *BnICS1_C* (C), *BnICS1_D* (D), and *BnICS1_E* (E). **(F)** Interactions of the CHE orthologs from *B. rapa* and *B. oleracea* with the Arabidopsis *ICS1* promoter. **(G)** Sequence alignment of CHE orthologs with different *ICS1* promoter-binding abilities. CHE orthologs lacking binding activity are highlighted in red. The red asterisk indicates the variable residue (glycine or lysine) within the TCP domain that correlates with binding capacity.

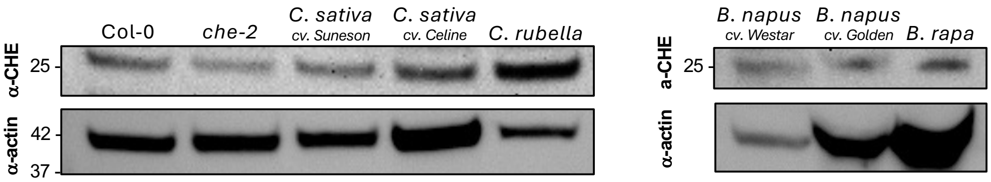


**Supplementary Figure 2. Immunoblot analysis using an anti-CHE polyclonal antibody.** Total protein extracts from plants used in the ChIP assay were subjected to immunoblotting with a polyclonal anti-CHE antibody. An anti-actin antibody was used as a loading control. Each CHE protein is expected to show a band at ~25 kDa. Actin shows a band at ~42 kDa. Protein samples were collected from Arabidopsis Col-0 and *che-2*, *C. sativa*, *C. rubella*, *B. napus*, and *B. rapa.*

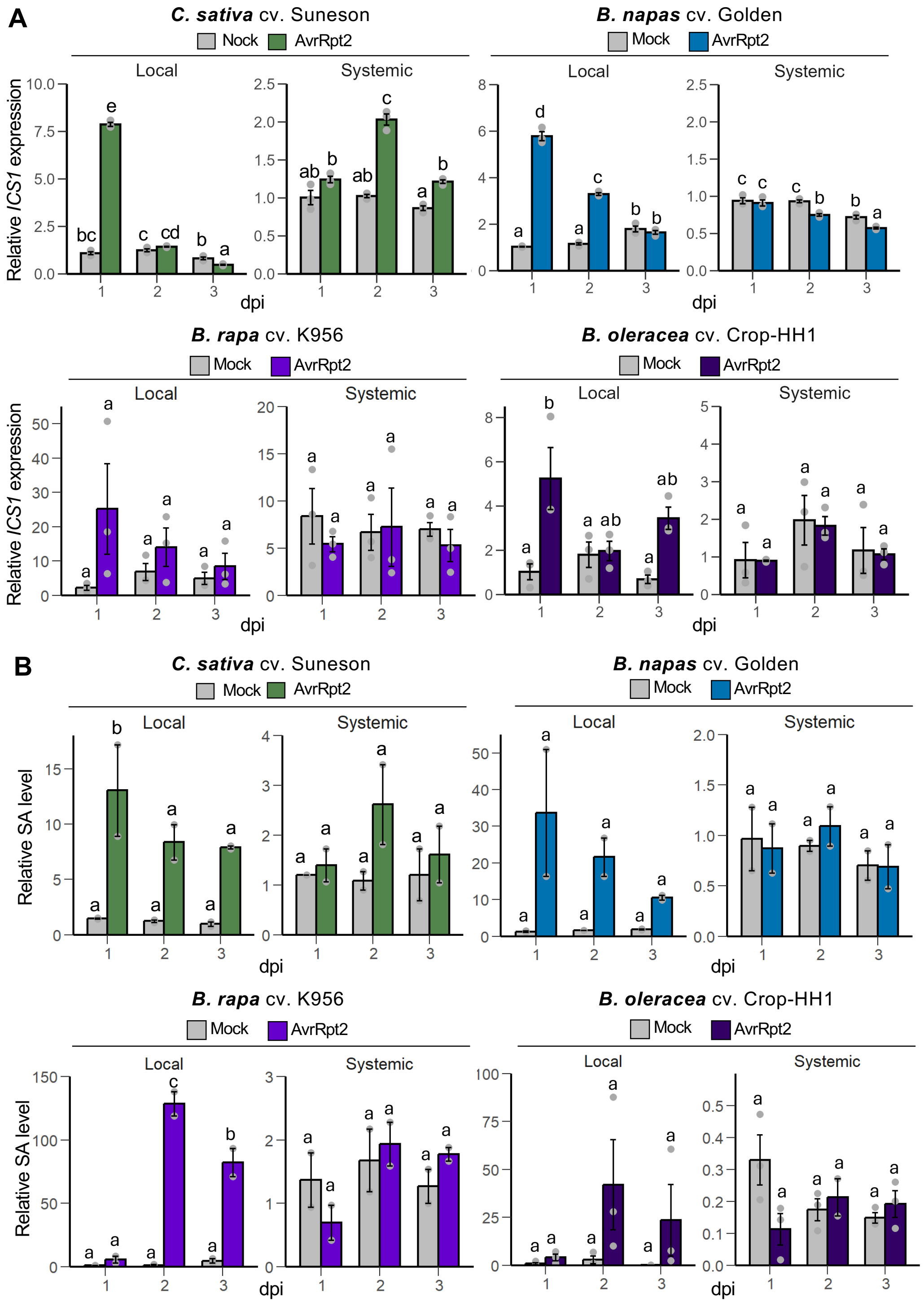


**Supplementary Figure 3. *ICS1* expression and SA accumulation in local and systemic tissue in Brassicaceae species.** Relative *ICS1* expression **(A)** and SA levels **(B)** in *C. sativa* cv. Suneson, *B. napus* cv. Golden, *B. rapa* cv. K956, and *B. olerace*a cv. Crops-HH1 following local infection with *Psm* ES4326/*AvrRpt2* or mock treatment. Different letters indicate statistically significant differences ($p < 0.05$, one-way ANOVA followed by Tukey's multiple comparisons test).
